# Neural code uses self-information principle to organize the brain’s universal cell-assembly coding

**DOI:** 10.1101/201301

**Authors:** Meng Li, Kun Xie, Hui Kuang, Jun Liu, Deheng Wang, Grace E. Fox, Zhifeng Shi, Liang Chen, Fang Zhao, Ying Mao, Joe Z. Tsien

**Affiliations:** Brain and Behavior Discovery Institute and Department of Neurology, Medical College of Georgia at Augusta University, Augusta, GA 30912, USA; The Brain Decoding Center, Banna Biomedical Research Institute, Yunnan Province Academy of Science and Technology, Xi-Shuang-Ban-Na Prefecture, Yunnan, China; Department of Neuropathology, Huashan Hospital, Fudan University, Shanghai 200040,China

**Keywords:** neural coding, cell assembly, neuronal variability, neuronal variability surprisal

## Abstract

The brain generates cognition and behavior through firing changes of its neurons, yet, with enormous firing variability, the organizing principle underlying real-time neural code remains unclear. Here, we test the Neural Self-Information Theory that neural code is constructed via the self-information principle under which each inter-spike-interval (ISI) is inherently self-tagged with discrete information based on its relation to ISI variability-probability distribution - higher-probability ISIs, which reflect the balanced excitation-inhibition ground state, convey minimal information, whereas lower-probability ISIs, which signify statistical surprisals, carry more information. Moreover, temporally coordinated ISI surprisals across neural cliques intrinsically give rise to real-time cell-assembly neural code. As a result, this self-information-based neural coding is uniquely intrinsic to the neurons themselves, with no need for outside observers to set any reference point to manually mark external or internal inputs. Applying this neural self-information concept, we devised an unbiased general decoding strategy and successfully uncovered 15 distinct cell-assembly patterns from multiple cortical and hippocampal circuits associated with different sleep cycles, earthquake, elevator-drop, foot-shock experiences, navigation or various actions in five-choice visual-discrimination operant-conditioning tasks. Detailed analyses of all 15 cell assemblies revealed that ~20% of the skewed ISI distribution tails were responsible for the emergence of robust cell-assembly codes, conforming to the Pareto Principle. These findings support the notion that neural coding is organized via the self-information principle to generate real-time information across brain regions, cognitive modalities, and behaviors.

## Introduction

A central theme in brain research is to understand how perception, memories, and actions are dynamically represented in real time by firing patterns of neurons. Over the past several decades, substantial efforts have been directed at measuring how changes in firing rates encode stimulus identity (referred to as the rate code), and/or how the exact timing of spikes from a neuron or the relative timing of spikes across multiple neurons convey information about the stimuli (generally referred to as the temporal code). However, one of the major stumbling blocks to cracking the *real-time* neural code is the neuronal variability; neurons in the brain discharge their spikes with tremendous variability across trials in response to the identical stimuli (1–5). From a signal-processing perspective, such a variability would represent noise, which has been shown to undermine reliable decoding of stimulus identities in real time (2, 3, 5–10). On the other hand, neurons in slice preparations are known to be capable of generating precisely-timed spikes in response to fluctuating currents injected at the soma (11, 12). Thus, firing variability is not due to imprecision in spike generation at the soma *per se*. Moreover, studies in the primary visual cortex or motor cortex showed that noise variability exhibited a certain level of correlation, which has been postulated to reflect varying attentional states or other modulatory signals such as intent (13–22). Despite the potential benefits or deeper implications that neuronal variability may bring, in practice, researchers typically treated firing variability as noise and dislodged it by applying over-trial data-averaging methods, e.g., peri-time stimulus histogram (PTSH) to better assess tuning properties of the recorded neurons. Obviously, such averaging procedure inherently assumed that any information possibly encoded in the temporal structure of the spike train can be (largely) ignored. Yet, it is widely acknowledged that this over-trial averaging approach is unlikely the real-time coding strategy that the brain uses to signal perception, memories, and actions on a moment-to-moment basis.

At a more fundamental level, firing variability - typically measured as variations in inter-spike-intervals (ISI) - is not just limited to trial-to-trial responses upon external stimulation, rather it represents a general property that is reflected continuously throughout all stages of mental processes and behaviors, including the “control” resting periods and sleep. In fact, spontaneous firing changes during those natural states or periods can be as large, if not larger, as those upon experimental stimulation (4, 8, 23–25). Such an ongoing spontaneous fluctuation in firing would seem to be profoundly paradoxical for achieving the robustness of neural coding on the moment-to-moment time basis: Why would the brain utilize this seemingly noisy and even counterproductive mode to convey information and generate cognitions in real time? Is there a deeper meaning for what neuronal variability may really stand? In other words, how is real-time neural code constructed in the face of such notorious firing variability? Can such a neural code be intrinsic to neurons themselves rather than being decipherable merely to outside observers?

In an attempt to explore these conceptual issues, we recently postulated the *Neural Self-Information Theory* that neural coding conforms to the self-information principle, which utilizes inter-spike-interval (ISI) variability and its variability history to generate the *self-information code* (26). Specifically, neuronal variability operates as the self-information generator and messenger; higher-probability ISIs, which reflect the balanced excitation-inhibition ground-state, convey less information, whereas lower-probability ISIs, which signify statistical surprisals, convey more information. At the physiological level, the rare-occurrence of ISI corresponds to unusually brief or prolonged neuronal silent periods, such as bursting patterns or strong inhibition, respectively. More importantly, the *Neural Self-Information Theory* further posits that at the population level, temporally coordinated ISI surprisals across neural clique members can seamlessly give rise to robust real-time cell-assembly codes (25, 26).

Guided by the *Neural Self-Information Theory*, we set out to test whether we can identify real-time cell assembly patterns from a variety of brain regions and across different mental states or tasks. Cell assembly is a group of co-activated neurons that has long been hypothesized by Hebb (1949) (27) as the population-level computational motif to represent real-time perception and thoughts (11, 28–32). We applied the *Neural Self Information Theory* to decode activity patterns in the prelimbic cortex, anterior cingulate cortex and hippocampal CA1 of freely-behaving mice. At least fifteen different cortical and hippocampal cell assemblies were identified in an unbiased manner, spanning from *those encoding categorical variables* such as earthquake, foot-shock, elevator-drop, or various operant actions during a five-choice visual-discrimination task, as well as those cell assemblies *encoding continuous variables*, such as spatial navigation or different stages of sleep cycles. Our analyses further revealed the conserved critical boundaries from which low-probability ISIs emerge as statistically significant self-information packets for the construction of robust cell-assembly neural code in real time.

## Results

### Neuronal variability is large and remains similar across the different brain regions

The *Neural Self-Information Theory* posits that neuronal variability acts as a self-information generator which should remain robust across various brain regions (26). On the other hand, if neuronal variability reflects *system noise*, one might expect that variability would grow larger as information is transmitted from low subcortical structures to high-cognition cortices. To differentiate these two scenarios, we used 128-channel tetrode arrays to record large numbers of neurons from six cortical and sub-cortical regions - namely, the basolateral amygdala (BLA), hippocampal CA1, dorsal striatum (STR), retrosplenial Cortex (RSC), prelimbic cortex (PRL) and anterior cingulate cortex (ACC), respectively - in freely-behaving mice. To facilitate direct comparison, we focused on putatively classified principal projection cells by excluding fast-spiking putative interneurons (Figure S1) and analyzed their variability distributions.

Neuronal variability is typically measured from ISIs distribution (Figure 1A) from which three well-defined statistics can be used to describe quantitatively neuronal variability of a given neuron - namely, a coefficient of variation (CV), skewness, and kurtosis (see Materials and Methods). In probability theory and statistics, CV is a standardized measure of dispersion of a probability distribution, and skewness is a measure of the asymmetry of a probability distribution, whereas kurtosis is a measure of the “tailedness” of a probability distribution. For comparison purposes, *normal distribution* (indicating stochastic process) has CV=mean/std, Skewness=0 and Kurtosis=0, while *exponential distribution* (indicating Poisson process) has CV=1, Skewness=2 and Kurtosis=6. The more skewed/long-tail a distribution exhibits, the larger CV, skewness and kurtosis it has.

**Figure 1.**
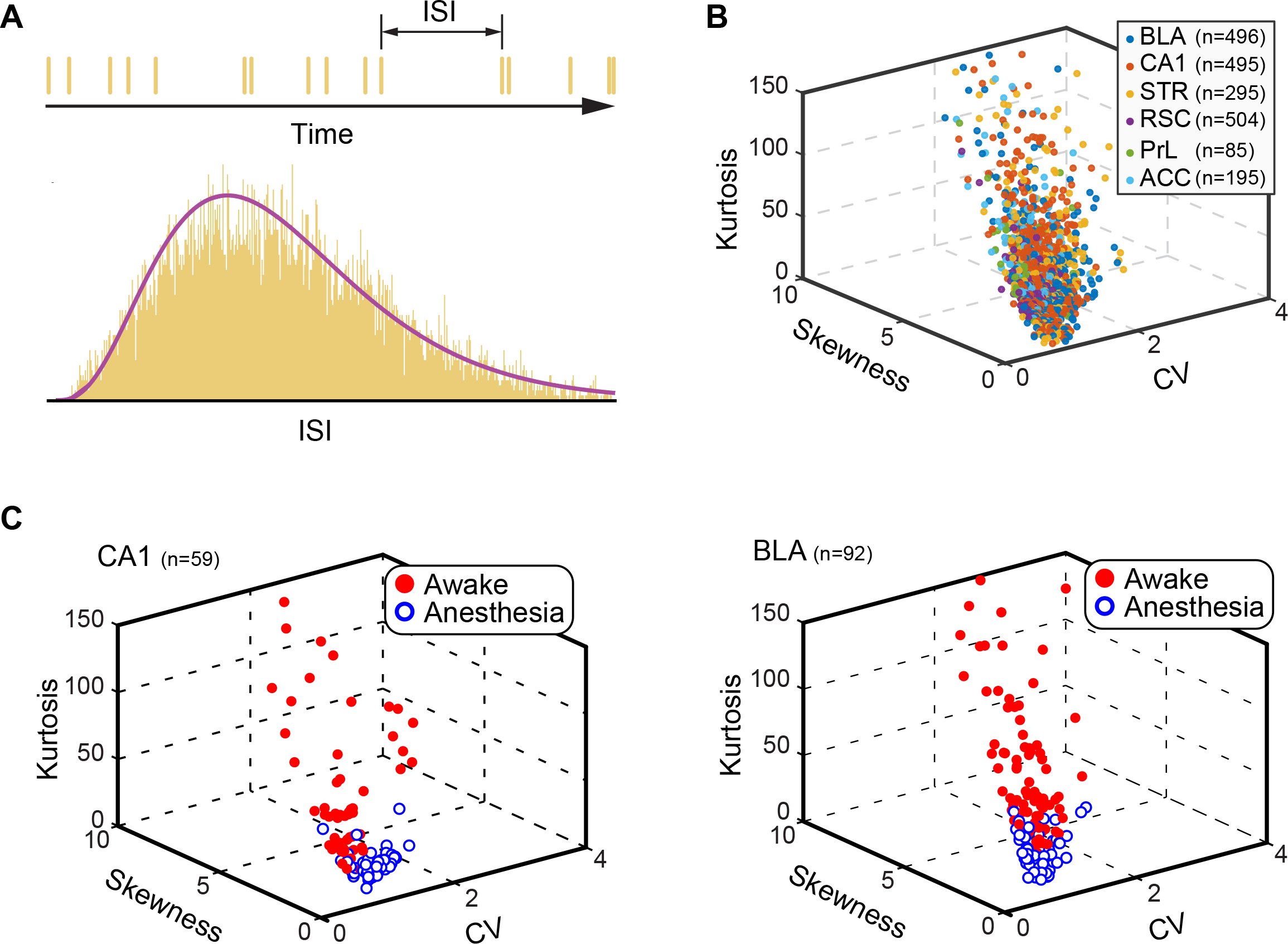
Neuronal variability remained similar across 6 different brain regions and was dramatically diminished upon anesthesia. (A) Neuronal variability can be assessed by inter-spike intervals (ISIs) distribution which follows long-tailed, skewed Gamma distribution. (B) The principle cells in six different brain regions show similar variability patterns. The basolateral amygdala (BLA), CA1, striatum (STR), retrosplenial cortex (RSC), prelimbic cortex (PRL), and anterior cingulate cortex (ACC). Numbers of principal cells from each brain region were indicated in the upper right box. Firing variability was assessed by Kurtosis, Skewness, and CV. (C) Neuronal variabilities were greatly reduced under anesthesia. Variability in the CA1 (left subpanel) and BLA (Right subpanel) was compared in using the same set of neurons recorded before (red circles) and after ketamine/dormiter injection (blue circles). Numbers of pyramidal cells used for analyses were listed on the upper left corner.

Our analyses showed that principal projection cells in these six different mouse brain regions exhibited similar neuronal variability distributions (Figure 1B, see also Figure S2A-D). Their CVs were all in a range of 0-4 (Figure S2B, BLA = 1.59 ± 0.02, CA1 = 1.64 ± 0.02, STR = 1.71 ± 0.03, RSC = 1.23 ± 0.01, PrL = 1.41 ± 0.04, and ACC = 1.54 ± 0.03). Skewness was in 0-10 range (Figure S2C, BLA = 3.27 ± 0.07, CA1 = 3.77 ± 0.07, STR = 3.74 ± 0.10, RSC = 3.00 ± 0.04, PrL = 3.47 ± 0.14, and ACC = 3.78 ± 0.11), whereas Kurtosis had a range of 0-150 (Figure S2D, BLA = 20.77 ± 0.94, CA1 = 29.45 ± 1.09, STR = 27.36 ± 1.47, RSC = 19.28 ± 0.66, PRL = 24.78 ± 2.04, and ACC = 28.98 ± 1.74). Therefore, these results show that neuronal variability is large and remains at a similar level across multiple brain regions and principal cell types (i.e. excitatory pyramidal cells in the CA1 vs. putative medium spiny neurons in the striatum). It did not grow larger from the amygdala and CA1 to the RSC and prefrontal cortices as the *system-noise model* would have predicted.

### Firing variability was reduced if cognitive coding was shut down by anesthesia

To further test the idea that neuronal variability serves as the self-information generator and messenger, one would expect that variability will diminish under the condition when both external and internal cognitive computation is artificially shut down (*i.e.* upon anesthesia). Ketamine/domitor administration is a widely used protocol to induced anesthesia. Accordingly, we recorded activity patterns of large numbers of cells from the BLA and CA1, respectively, in the awake state as well as under ketamine/domitor-induced anesthesia. Using the same set of putative pyramidal neurons, we asked how the skewed distributions of ISI differ during anesthesia vs. those during awake period. Indeed, in both the BLA and CA1, putative pyramidal cells drastically decreased their neuronal variability upon anesthesia (Figure 1C; also see Figure S2E). The skewed/longtailed distributions of ISI in the CA1 was significantly reduced (CV: 1.71 ± 0.08 in awake vs. 1.36 ± 0.21 under anesthesia, ***P***=0.00024; skewness: 4.46 ± 0.25 in awake vs. 2.04 ± 0. 38 under anesthesia, ***P***=4.1e-16; Kurtosis: 39.53 ± 4.17 in awake vs. 8.68 ± 2.85 under anesthesia, ***P***=3.3e-11) (Figure S2E, top row). Similarly, BLA pyramidal cells also greatly reduced their neuronal variability (CV: 1.83 ± 0.05 in awake vs 1.28 ± 0.20 under anesthesia, *P*=1.1e-16; skewness: 4.90 ± 0.19 in awake vs. 2.46 ± 0.50 under anesthesia, ***P***=5.8e-24; kurtosis: 45.29 ± 3.54 in awake vs. 11.87 ± 3.92 under anesthesia, ***P***=4.0e-17) (Figure S2E, bottom row). The reduction in firing variability was evident from ketamine-induced rhythmic spike-discharge patterns (Figures S2F and S2G). These pharmacological intervention experiments demonstrate that the shutting down of external and internal cognitive coding processes indeed diminished neuronal variability.

### *Self-Information theory-based* decoding strategy to uncover cell assemblies

The key feature of the *Neural Self-Information Theory* is that any given ISI is inherently self-tagged with a discrete amount of information based on its ISI (the silence time duration) in a relationship with the ISI variability-probability distribution (variation history of silence time durations). Specifically, the amount of self-information (“*SI*”) contained in each ISI can be quantitatively obtained based on the ISI variability-distribution probability [*SI* = - log(*p*), where *p* is the probability] (26). The smaller the probabilities in ISIs, the larger the variability surprisals, which convey more information (in statistical terms, those events with low-occurrence probability are called *surprisals*). At the physiological level, these self-information ISI surprisals can be either positive (when a neuron’s ISI becomes much shorter than is typical, reflecting strong excitation) or negative (when ISI becomes much longer than is typical, reflecting strong inhibition). Subsequently, these dynamic, transient surprisal silence-duration (ISI) patterns would act as the critical realtime information packets at a single neuron level. More importantly, when these surprisal ISIs are generated across a cell population in a temporally coordinated manner, their collective patterns would intrinsically give rise to robust real-time cell-assembly code.

To experimentally test the validity of this hypothesis, we set out to devise a general decoding strategy based on the neural self-information concept to uncover cell assemblies from spike-train datasets recorded from different brain regions, mental states, and behavioral tasks. This <u>V</u>ariability-based <u>C</u>ell-<u>A</u>ssembly <u>D</u>ecoding (VCAD) method consisted of the following three major steps (Figure 2):

**Figure 2.**
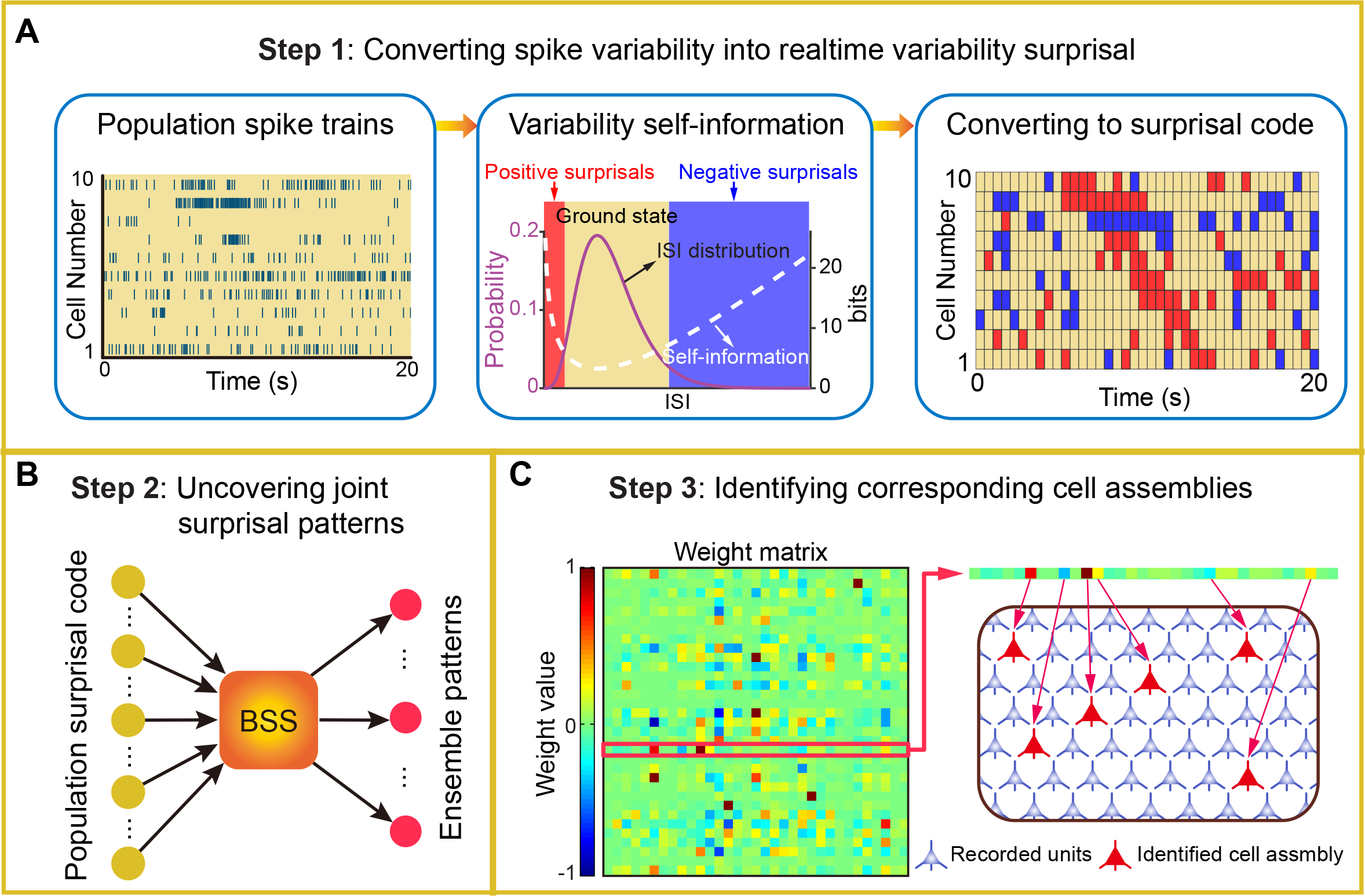
The strategy to apply the Self-Information Coding concept to decode cell assembly patterns in an unbiased manner. (A) Step 1: Converting spike variability into real-time variability surprisal. The spike activities of 10 simultaneously-recorded cells are illustrated in the left subpanel, the ISIs of each cell is fitted by the Gamma distribution model which assigns each ISI with a probability. Based on the fitted Gamma distribution model, two boundaries are then assessed (as shown in the middle subplot) to generate three states; that is, positive-surprisal state (short ISIs), negative-surprisal state (long ISIs), and low self-information ground state. As such, neuron’s spike trains can be converted into ternary surprisal codes [excitation surprisal as 1 (red blocks), ground state as 0 (yellow blocks) and inhibition surprisal as-1 (blue blocks)]. (B) Step 2: Uncovering joint surprisal patterns. Taking population ternary surprisal code as input, the ensemble patterns were unbiasedly discovered by the blind source separation (BSS) method which aims for searching joint variability-surprisals in both space (across simultaneously-recorded cells) and time (moment-to-moment dynamics). (C) Step 3: Identifying corresponding cell assemblies. For each decoded independent signal source, neurons’ information contributions are scaled quantitatively by the absolute values in the weight matrix *W* of BSS analysis (shown in the left subpanel), thus, corresponding cell assemblies can be identified by picking up top-weight cells (right subpanel).

#### 1) To convert ISI variations into real-time variability surprisals

In practice, measurement of ISI probability distribution can be determined numerically. In contrast to the prevalent notion that spike variability is a Poisson distribution, emerging studies have suggested that ISI variability in many neural circuits conform to the gamma distribution (33). Thus, we first fitted single neuron’s ISIs with a gamma distribution model which can assign each neuron’s ISI with a probability. Subsequently, a spike train emitted by a neuron can be transformed into surprisal-based ternary code (positive surprisal as 1, ground state as 0, negative surprisal as -1) to describe the dynamic evolution in self-information states (Figure 2A).

#### 2) To uncover joint ISI surprisal patterns in space and time

This step searches for joint variability surprisals in both space (across simultaneously-recorded cells) and time (moment-to-moment dynamics). Blind source separation (BSS) methods, such as independent component analysis (ICA), can be used to efficiently identify a set of independent information sources from simultaneously observed signals as structured patterns or relationships. Although the nature of the demixing matrix’s dimension *p* meant that *p* information sources can be theoretically decoded from population activity, we reasoned that optimal neural coding should also be energy efficient via utilizing the least amount of variability surprisals together with the minimal numbers of such information-coding cells. As such, we used the minimal CV values in each dataset to unbiasedly assess the optimal numbers of independent information sources (distinct cell assemblies) (Figure 2B; also see Figure S3A, Materials and Methods).

#### 3) To identify corresponding cell assemblies

Each independent signal source decoded by BSS corresponds to a distinct real-time activation pattern given by a cell assembly. To discover the functional meaning of each cell assembly pattern, one can compare the time points marked by each real-time activation with various other experimental parameters recorded during experiments, such as local field potential (LFP), the time points of stimulus presentations, or video tapes of an animal’s behavioral state, specific actions and performance, or spatial locations, etc. The top-ranking membership with the highest contribution weights in the cell assembly can be directly identified from demixing matrix **W** (Figure 2C; also see Figure S3B). Moreover, their contributions to a given cell-assembly pattern can be quantitatively defined by shuffling techniques (i.e. by shuffling or artificially changing spike patterns to alter surprisal states). This step allowed us to assess quantitative membership information that other dimensionality-reduction-based pattern-classification methods (i.e. principal component analysis or multiple discriminant analysis) could not provide.

In the study of neural coding, cognitive and physiological inputs associated with external and internal states typically fall into two major categories - namely, continuous variables (i.e. arm movement, navigation, sleep oscillations, etc.) and categorical variables (i.e. distinct events, stimuli, discrimination tasks, etc.). If neuronal variability surprisals act as the universal coding vehicle to provide discrete quanta of information, one would predict that this VCAD method should be able to uncover various cell assemblies related to such a wide range of dynamic operations across different brain regions.

### Identification of cortical cell assemblies encoding distinct emotional experiences

We asked whether a variability-based self-information process underlies real-time coding of discrete categorical variables, namely, fearful experiences such as earthquake, foot-shock and a sudden elevator drop (as one experienced during the Tower-of-Terror ride in Disney theme park). The anterior cingulate cortex (ACC) is part of the prefrontal cortex known to process emotions and fear memories (34–36). To examine how the ACC discriminates and categorizes distinct fearful experiences at the population level, we employed 128-channel tetrodes to monitor spike activity of large numbers of ACC while the recording mice encountered earthquake, foot-shock and sudden elevator drop which are known to produce fear responses (37). By scanning through the real-time spike dataset that contained 146 well-isolated, simultaneously-recorded ACC units, our VCAD method automatically uncovered three distinct ensemble patterns. We found that these patterns corresponded to the occurrences of one of three fearful stimulations (Figure 3A). Specifically, ACC assembly-1 activations were time-locked to the occurrences of six earthquake events; ACC assembly-2 was temporally corresponding to six foot-shock events; and ACC assembly-3 was matched to free-fall events. Their unique ensemble patterns were further verified by shuffling their top 20% large-weight neurons’ firing patterns with the Gaussian signal using the same mean firing rate and standard deviation (Figure S4A). This shuffling procedure showed that the ensemble representation of a given event (e.g. free-fall) gradually became weaker as more top-contribution neurons were shuffled, while leaving the other two ensemble patterns (e.g. representing the earthquake or foot-shock) unchanged. Peri-event spike raster and histogram plot analyses (Figure S4B) revealed that while the majority of member cells exhibited event-specific responses (i.e., earthquake-specific, free-fall specific, or foot-shock specific ACC cells), some of the top cell-assembly members participated in two cell assemblies or even all of the three assemblies (Figures 3B-D).

**Figure 3.**
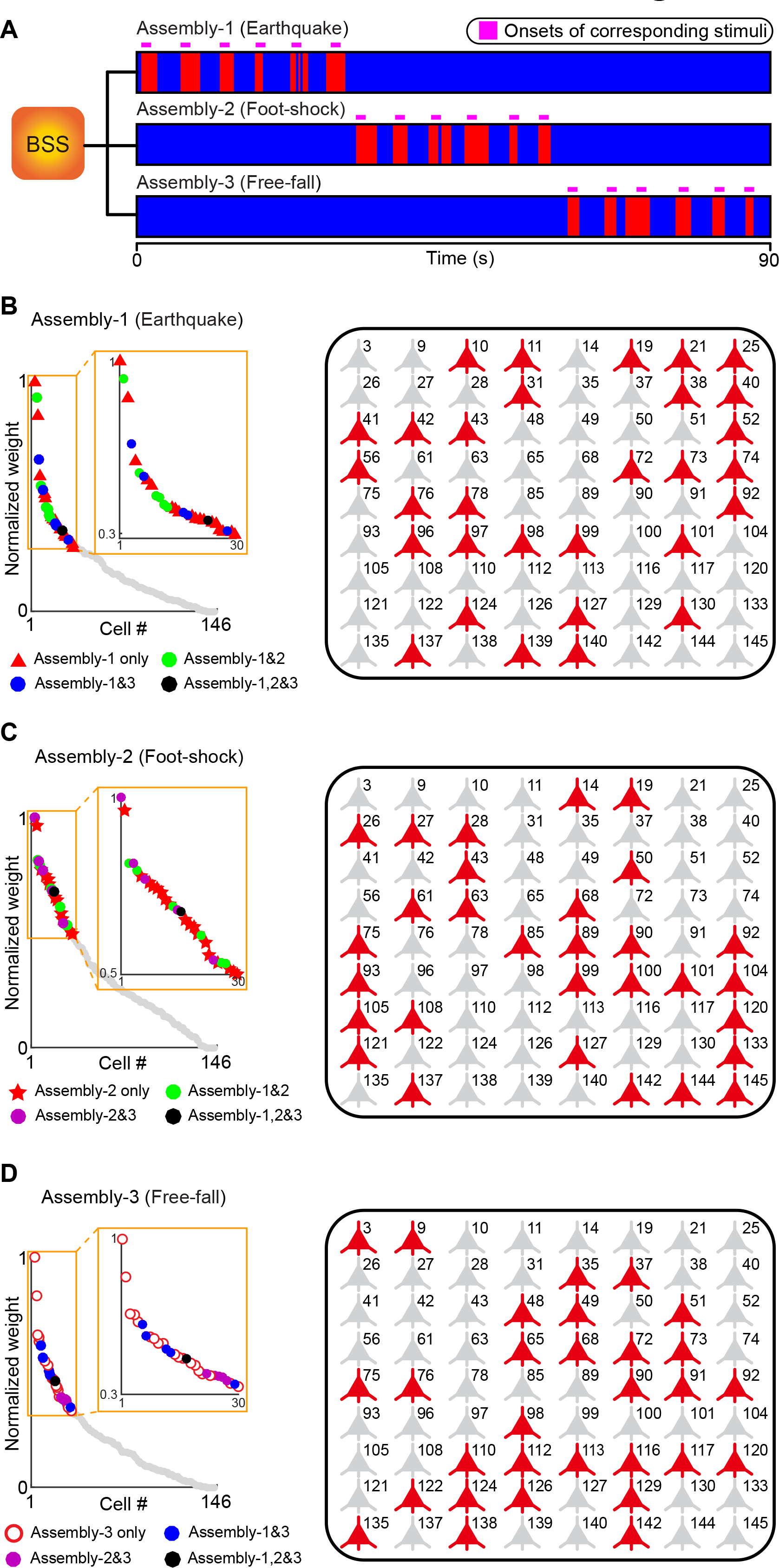
Identification of three ACC cell assemblies 1007 corresponding to three distinct fearful experiences. (A) Three cell assemblies are decoded from ACC during fearful startle experiment. Each cell assembly corresponds to the occurrences of one of three fearful startle events (Six trials per event, as shown by six purple bars above each ensemble patterns). (B) Left panel: Cell membership analyses of Assembly-1 show that some neurons participate in more than one cell assemblies, using a specific-to-general combinatorial coding manner. Right panel: illustration of identified Assembly-1 member cells engaged in representing Earthquake (in red color). Non-member cells are in gray. The numerical number on the top-right corner of each cell represents the unit simultaneously recorded, arranged from the 128-channel array. (C) Assembly-2 corresponding to Foot-shock. (D) Assembly-3 corresponding to Free-fall.

### Identification of cell assemblies related to sleep oscillations

Next, we investigated whether the self-information coding concept can be used to identify cell-assembly patterns related to internally-driven, continuous variable processes. Sleep is associated with and defined by changes in EEG or local field potential oscillations (38). It has important functions for memory consolidation and behaviors (38–41) or fear-memory consolidation (42–44), relatively less is known about the functional classification and organization of CA1 cell assemblies during natural sleep oscillation cycles. Thus, we asked whether the VCAD method can be used to identify cell assemblies in the CA1 related to various cycles of sleep. We scanned a 20-minute spike dataset collected from the mouse hippocampal CA1 region during a sleep session and analyzed the ISI variability of 266 units recorded simultaneously using 128-channel tetrode arrays. Our VCAD analyses of spike dynamics unveiled a total of three major ensemble patterns over the time course of 20-minute sleep period (Figure 4A). To search for the relationships of these three assembly patterns and sleep oscillations, we aligned these CA1 temporal patterns with the time-course plot of the simultaneously recorded LFP oscillations. Interestingly, we found that the temporal emergence of Cell Assembly-1 patterns was tightly matched to the occurrences of theta oscillation (Figure 4B), whereas Assembly-2 patterns were time-locked to ripple oscillation (Figure 4C). Furthermore, we found that the activation of Assembly-3 temporally corresponded to the DOWN-state of sleep cycles (Figure 4D) - that is, this ensemble pattern was consistently time-locked with the troughs of various LFP frequency bands.

**Figure 4.**
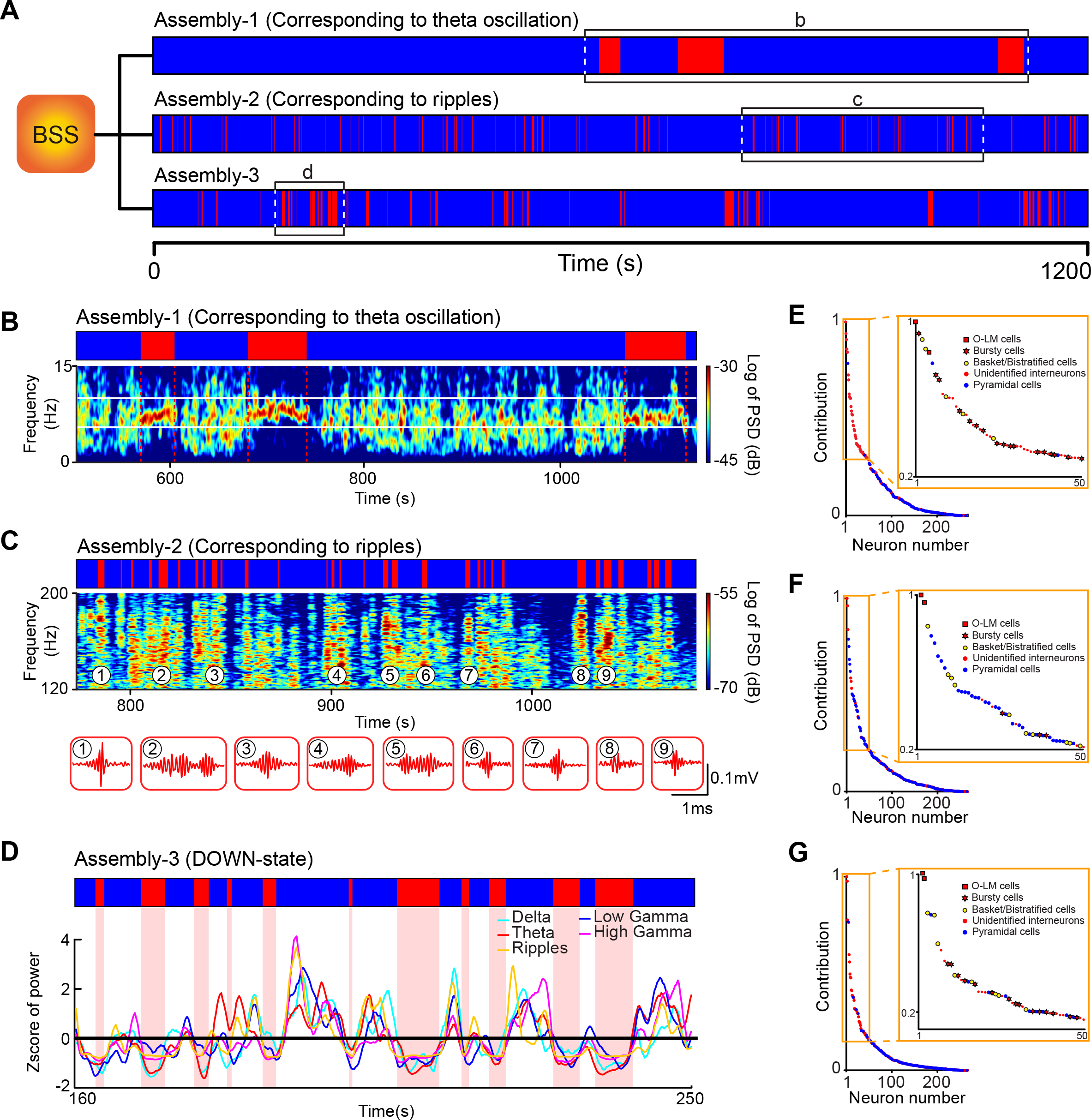
Three distinct CA1 cell assemblies identified from the different sleep cycles. (A) Three information sources (ISs)-based ensemble patterns are decoded by VCAD method during animal’s sleep. (B) Assembly-1 (Corresponding to theta oscillation stage). The upper bar shows the activities of decoded theta ensemble pattern, the red blocks denote the activations of Assembly-1, while the blue blocks represent its inactivity. The power spectrogram below shows the simultaneously-recorded local field potential activity within 0-15 Hz frequency band, the theta band (6-10 Hz) is highlighted by white lines. The decoded ensemble pattern is in complete correspondence with the elevated theta band activity (as shown by the red dashed lines). (C) Assembly-2 (corresponding to ripples). (D) Assembly-3 (corresponding to DOWN-state). Z-scores of power for well-defined frequency bands (Delta, Theta, Low Gamma, High Gamma, and Ripples), where the black line (Z-score = 0) denotes the mean power in each band. It can be observed that the ensemble pattern of Assembly-3 is consistently time-locked with the troughs of these LFP frequency bands. (E) Analyses of different cell types consist in Assembly-1. Most cell members of Assembly-1 are putative interneurons. The ripple-band (120-200 Hz) power spectrogram is shown in lower subpanel for verifying the accuracy of cell assembly decoding result. Nine ripple waveforms are replotted below. (F) Cell-type analyses show that top Assembly-2 cells were mostly putative pyramidal cells. (G) Cell-type analyses of Assembly-3 also shows that its members were mostly interneurons.

By further taking advantage of the VCAD method in providing the ranking of cell-assembly members, we found that the top 20% ranking units (with largest weights in demixing matrix **W**) in CA1 Assembly-1 cells with consisted of mostly putative interneurons (Figure 4E, n=53 cells). The shuffling technique (replacing their firing pattern with Gaussian signal with the same mean firing rate and standard deviation) revealed that the Assembly-1 pattern was abolished as these top 20% contribution-cells’ firing patterns were shuffled (Figure S5A). Interestingly, only about 15% of these Assembly-1 cells (eight cells, identified as putative O-LM cells or basket cells/bistratified cells based on the multiple criteria for interneuron sub-classification, see Figure S5B and S5C) exhibited a robust phase-locking relationship with theta oscillations (termed as Theta-coupled Assembly-1 cells) (Figure S5B and S5C), which is consistent with evidence from literature that CA1 interneurons exhibited theta-coupling (39, 40, 45–47). Surprisingly, many Assembly-1 cells (45 cells, ~85% in Assembly-1) were not phase-locked with theta phase, but rather exhibited a significant difference in averaged firing rates between theta epochs vs. non-theta epochs. One major group of the rate-altering cells, classified as putative Bursty cells (17 cells) (Figure S5D), dramatically increased their firings during theta cycles in comparison to non-theta epoch, whereas two putative pyramidal cells significantly decreased their firings during theta sleep cycles (Figure S5E).

Another surprizing finding was that many theta-coupled CA1 cells from the simultaneously recorded dataset did not belong to Assembly-1. These non-member theta cells were intriguingly distinct from those Assembly-1 theta cells as confirmed by their significant differences in firing correlation (Figure S6A). This suggests that theta-coupling does not necessarily indicate the common membership. In fact, two distinct populations of theta interneurons exist in the CA1, with one subset uniquely engaged in sleep theta. Similarly, we noted that there were many rate-altering cells that did not belong to Assembly-1. Their distinct memberships were further supported by a dramatic statistical difference in the correlation coefficient between rate-altering member cells and rate-altering non-member cells (Figure S6B).

We then analyzed the nature of CA1 Assembly-2 ensemble patterns which were tightly time-locked with ripple peaks (Figure 4C). In contrast to Assembly-1, the majority of the top Assembly-2 cells were mostly putative pyramidal cells (Figure 4F). It is worthwhile to note that many *non-member cells* also showed rate changes between ripple epochs vs. non-ripple periods. Yet, they were distinct from Assembly-2 cells as evidenced by the significant difference in their correlation (Figure S6C).

As to Assembly-3 cells (Figure 4D), we found that they also consisted of mostly interneurons (Figure 4G). But many of them were different from those interneurons listed in the Assembly-1 (Figure S7). It was repeatedly noticed that several cells (~15%) were cross-listed in multiple-sleep oscillatory patterns. The best example is that the same putative O-LM cells were repeatedly identified in all three sleep assemblies. Overall, these three distinct cell assemblies identified by the self-information coding scheme consistently showed significantly higher correlations among themselves, in comparison to across the populations (Figure S6D). The above results demonstrated that the unbiased VCAD method has enabled us to identify distinct cell assemblies that were time-locked to distinct sleep LFP oscillations.

### Unbiased identification of CA1 cell assemblies underlying place navigation and start/finish experiences

To further demonstrate the generality of the self-information coding concept, we asked whether the VCAD method can be used to identify and verify the most studied cell assembly patterns, namely, place cells in the hippocampus. CA1 place cells exhibit sequential firing patterns indicating positional information (23, 28, 30, 31, 48–54). Similar to the huge variability of other neurons in the brain, place cells also exhibit trial-to-trial variability, which rendered real-time firing sequences unreliably (23, 24, 28, 30, 31). Traditionally, identification of place cells involved averaging spike counts over spatial tiles using x-y pixel coordinates. Because motion can greatly change firing rates of CA1 cells, in literature spike rasters below a certain running speed were typically removed during this data analysis step (a process critical to dislodge variability noise). We wondered whether the self-information concept could be used to identify hippocampal place cells without using such trial-averaging methods, but rather in an unbiased manner via simply scanning population spike rasters using the VCAD sliding-window method on a moment-to-moment fashion.

Therefore, we applied the self-information decoding method to 20-minute spike trains of 266 simultaneously recorded CA1 units from a mouse which was well trained to run back and forth on a one-meter linear track (without using trial-averaging spike counts over spatial pixels). Based on the minimum CV as the best relevant dimension, our VCAD method unbiasedly revealed two distinct CA1 cell ensemble patterns (Figure 5A). As we aligned these two cell-assembly patterns with a videotape that recorded the animal’s navigational positions and behaviors on the linear track, we found that these two real-time ensemble patterns were matched nicely to the westbound and eastbound navigations, respectively (Figure 5B). Then, we examined top-ranking members from each cell assembly based on their matrix weights, and performed position-firing analysis of these cells. A closer examination of the top-weighted CA1 neurons listed in these two cell assemblies revealed that some of these cells exhibited classic single place field place (Figure 5C). Many of other cells also showed multiple place fields (Figure 5D). Among the list, we also noticed that some of the top-contributing CA1 cells did not exhibit discrete place field(s), but rather had differential responses corresponding to directional routing as the mouse traveled westbound or eastbound (Figure 5E) - that is, some CA1 cells preferentially fired in one travel direction over the other.

**Figure 5.**
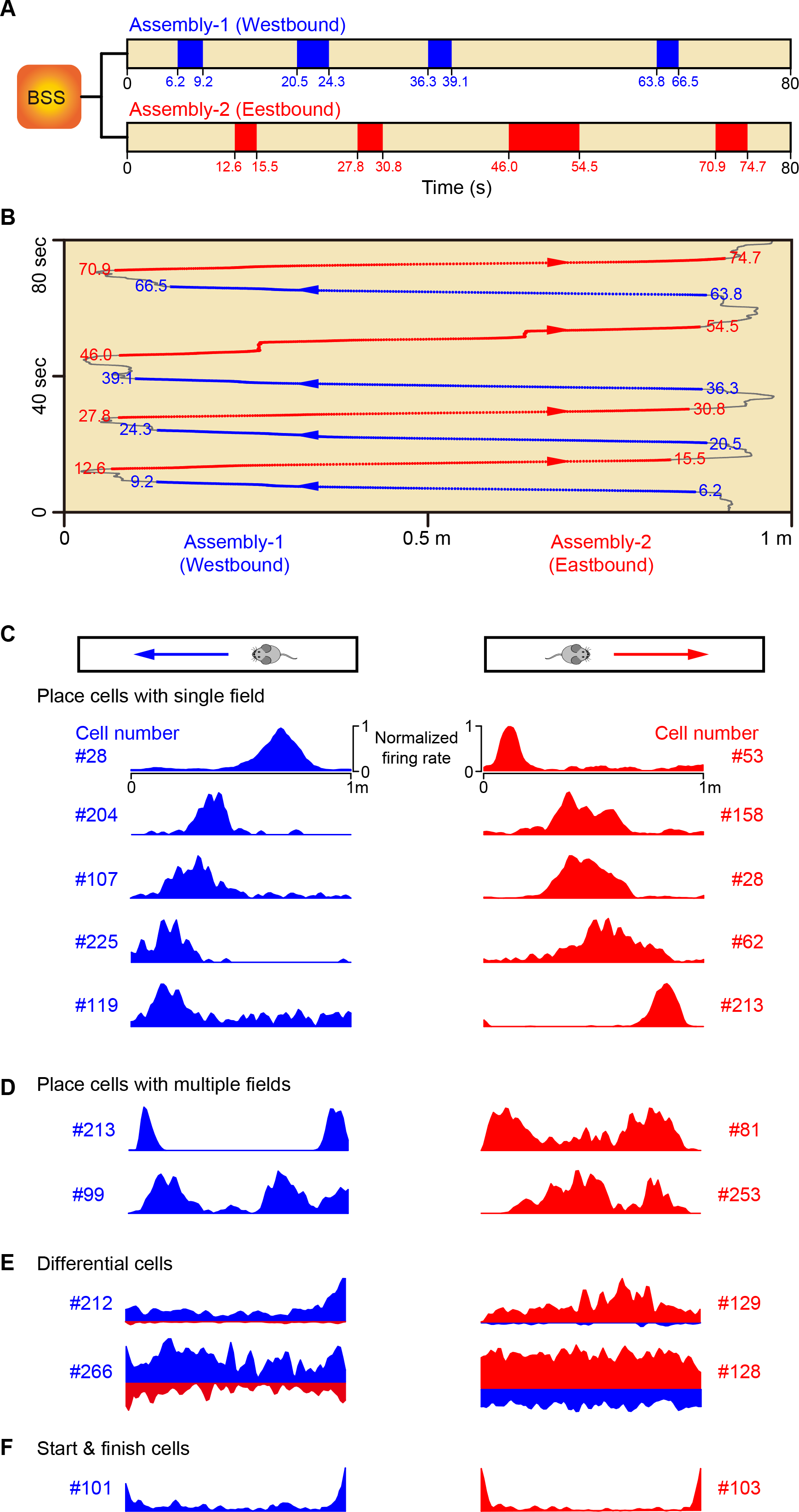
Unbiased identification of two navigation-related CA1 cell assemblies during linear track exploration. (A) Two information sources (ISs)-based ensemble patterns were decoded by VCAD method. The blue/red blocks denote the activations of each corresponding ensemble patterns. (B) The animal’s trajectory within 80s during the linear track experiment (1-m length). Blue dots and arrows denote the decoded ensemble pattern corresponding to “Westbound,” while red dots and arrows are for “Eastbound.” (C-F) Firing patterns of top-weight member cells in two cell assemblies. These top-weight cells fall into four categories: place cells with single place field (C), place cells with two place fields (D), differential cells (E) which exhibited significantly higher firing rates in one of the directions than the opposite direction, P<0.001 through pairwise t-test). (F), Start & finish cells that suppressed their firings during the navigation. Two cells exhibiting westbound-or eastbound-specificity are shown.

Unexpectedly, from the list of the VCAD-identified cell assemblies, we also found that both the eastbound and westbound cell assemblies contained cells signaling start/finish transitions, that is, these cells specifically decreased their firing at the start of running and remained low-firing until the end of the navigation (Figure 5F). While we found two start/finish cells engaged in both the westbound and eastbound journals, most interestingly, several CA1 start/finish cells exhibited journal-or direction-specific firing decreases (Figure 5F Unit-101 in the left sub-plot vs. Unit-103 in the right sub-plot; also see Figure S8). Such route-specific start/finish responses strongly suggest that the information coded by these cells were not merely corresponding to changes in motion states such as speed acceleration/deceleration, but journey-specific episode(s). Therefore, the neural self-information coding concept has successfully allowed uncovering CA1 place cells and their other assembly members, such as start/finish cells, important to account for multiple aspects of navigation-related variables.

### Prefrontal cell assemblies encoding five-choice visual-discrimination operant-conditioning tasks

Finally, we applied the self-information decoding approach to uncover novel cell assemblies in the prefrontal cortex related to the five-choice visual discrimination operant conditioning task – one of the most classical behaviors requiring a set of visually- and spatially-guided procedural actions. In this task, mice learned to nose-poke in one of the five temporarily lit apertures within a short time window in order to receive a food reward delivered from the food magazine located on the opposite side of the chamber. The mice were trained to reach an 80% success rate (see Materials and Methods). A successful trial typically included the 5-sec illumination of one of the five apertures (in a randomized fashion) to signal the beginning of the trial. During this time, the mouse needs to run from the food magazine area to the response aperture area and then nose-poked in the transiently lit aperture (correct responses), which resulted in a single food pellet dropped onto the food dish from the magazine. The mouse then runs back to the food magazine area to eat the pellet, which triggers the next trial. A single session typically consisted of 50 trials or more during which mice performed this attentive choice-action task.

We used 128-channel tetrode arrays and recorded from the prelimbic cortex (PRL) while well-trained mice performed this operant conditioning task. A total of 100 PRL cells were identified as a well-isolated unit and used for present analysis (those units did not meet the criterion were excluded). We then scanned the spike dataset consisting of these PRL units using the VCAD method, and a total of seven cell-assembly patterns were unbiasedly detected. We then align the temporal occurrences of these seven distinct patterns with the recorded videotape, and found that all of them could be temporally matched to distinct stages of the operant procedural task (Figure 6A, the sequential ensemble patterns from three back-to-back trials were plotted).

**Figure 6.**
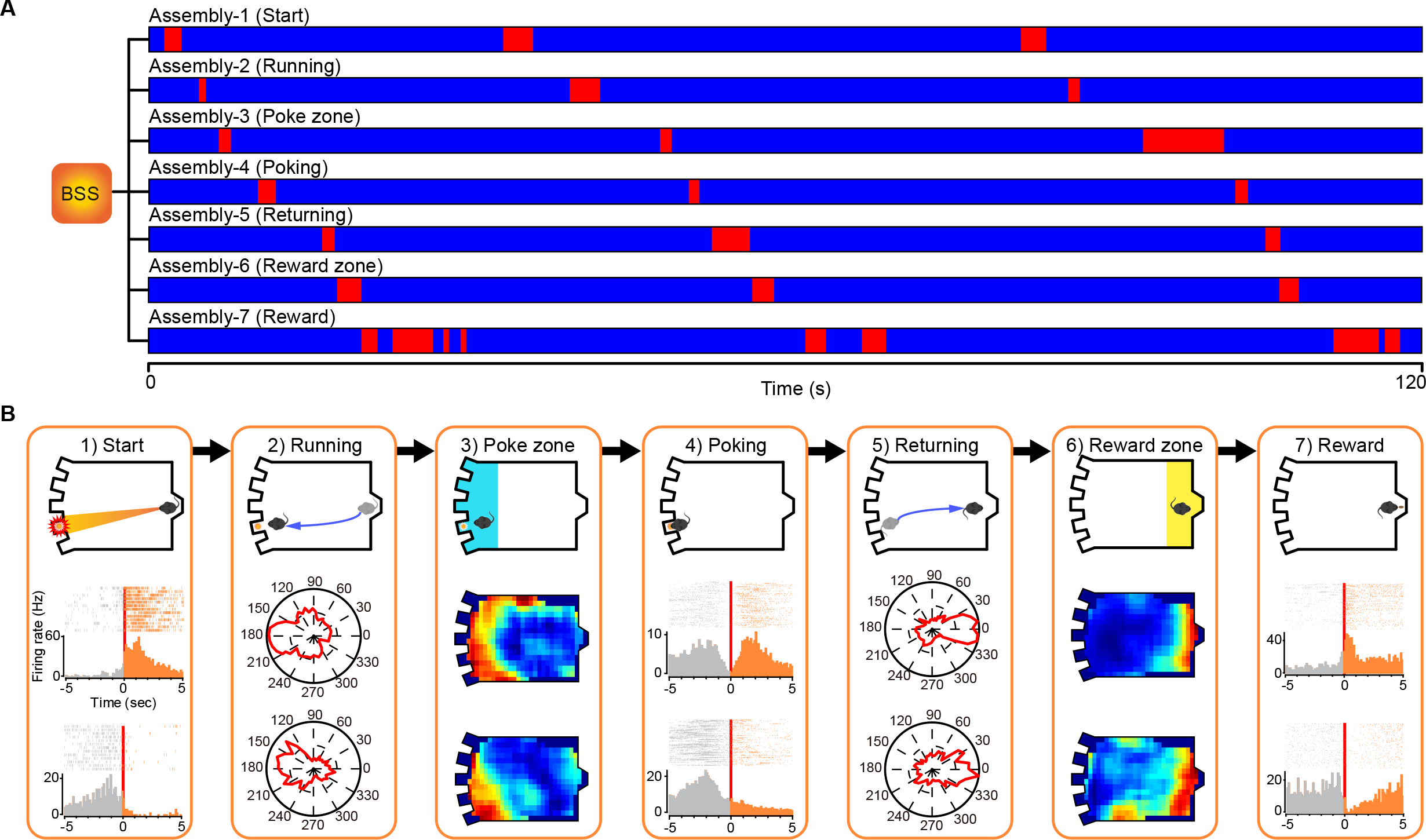
Seven PrL cell assemblies were identified during five-choice discrimination operant-conditioning task. (A) Activations of seven decoded ensemble patterns during three back-to-back trials of five-choice discrimination operant-conditioning task, each ensemble pattern corresponds to one of seven distinct stages of the operant-procedural task. (B) Illustrations of seven decoded ensemble patterns and examples of top large-weight cells in each cell assembly. **Stage-1 (Start)** corresponds to the moment which visual stimulus (the illumination of the aperture light on) was presented in one of five spatial locations to start a trial. Peri-event rasters/histograms of two example cells exhibited significant increased\decreased firing activities upon the illumination of the aperture light (time = 0). These two cells were anticorrelated, the correlation coefficient between these two cells was -0.337. **Stage-2 (Running)** corresponds to the route when the animal runs from food magazine region to response aperture region. Orientation-firing polar histograms of two large-weight cells show obvious orientational preferences. **Stage-3 (Poke zone)** activated when the animal was near the response apertures. Apparent spatial-specificity firing patterns of top-contributed cells were observed when the place-cell analysis was applied. **Stage-4 (Poking)** corresponds to the poking of the response aperture. Obvious firing pattern changes were observed upon the poking (time =0) in the peri-event rasters/histograms of top-contributed cells. **Stage-5 (Returning)** activates when the animal was running back from the response apertures to the food magazine. Obvious orientational preferences of cell-assembly members were observed in the orientation-firing-polar-histogram analyses. **Stage-6 (Reward zone)** corresponds to the locations near food magazine. Place cell analyses verify that the cell-assembly members exhibited higher firing probabilities within the location near the food magazine. **Stage-7 (Reward)** corresponds to the actions of collecting the reward pellet. Increased/decreased firing activities of two cell-assembly members are shown in the peri-eventrasters/histograms.

The PRL assembly-1 pattern was time-locked to the cue-onset when the stimulus aperture was transiently lit for 5-second, signaling the initiation of the five-choice visual discrimination task (Figures 6A and 6B). The peri-event spike raster and histogram confirmed that Assembly-1 cells exhibited significant changes upon the aperture *light on* (Figure 6B, Stage-1). For example, A top-ranking PRL unit increased its firing upon the light-on in the stimulus aperture (5-second duration). The firing of this attentive cell was then tapered off gradually (Figure 6B, listed in 1) Start. Upper PTSH subplots). Another PRL cell suppressed its firing upon the stimulus aperture light-on (Figure 6B, 1) Start. Bottom PTSH subplots).

The PRL assembly-2 activation pattern was time-locked to the first cue-conditioned operant action, namely, running towards the light-lit stimulus aperture from the reward zone toward where the animal was originally located (Figure 6B, Stage-2; two PRL cells are shown). The orientation-firing polar histograms of comparing these cells’ activity patterns during the five-choice visual discrimination operant task with those during home-cage running behavior showed that these firings were specific to the operant-conditioning task (Figure S9A).

The PRL assembly-3 pattern corresponded to the entering of the poke zone (the region near apertures) (Figure 6B, Stage-3). The place-firing analysis method confirmed that these large-weight responsive PRL units exhibited poke zone-specific spatial firing patterns (two cells were listed in the upper and lower subpanels, respectively).

Firing patterns of the PRL assembly-4 cells were time-centered around the preparation and the actions of nose-poking at the transiently lit apertures (Figure 6B, Stage-4). Peri-event spike histogram analysis revealed that these units exhibited dynamic firing changes around the nose-poking action. One representative PRL cell exhibited transient suppression during poking of the stimulus aperture (the upper PTSH subpanels), whereas another cell peaked its firing during the preparation phase of nose-poking (lower subpanels), consistent with the role of prefrontal cells in action planning of goal-execution.

The PRL assembly-5 pattern corresponded to the 4^th^ action phase, namely, running back from the poke zone toward the reward zone where the food magazine was located (Figure 6B, Stage-5). The orientation-firing polar histograms revealed elevated firing changes by this type of member cells. Again, these PRL cells did not show any significant firing changes when the mouse was running around in its home-cage environment (Figure S9B), suggesting that these cells were goal-oriented.

Increased firings of the PRL Assembly-6 cells were time-matched to the animal’s arrival in the reward zone (Figure 6B, Stage-6, two example cells were listed). These cells’ location-specific firing changes were evident from the PTSH plots again their spatial positions at the reward zone.

Finally, the Assembly-7 cells decoded by the VCAD method responded to consumption of food pellets. These PRL cells either increased or decreased their firings (Figure 6B, Stage-7, two cells were shown). While many of the above cells tended to be specific to a given phase of the five-choice operant-conditioning task, we noted that some of the cells participated in multiple cells assemblies. For example, some PRL cells belonged to both the running action assembly and stimulus-aperture zone assembly (Figure S9C). other PRL cells would exhibit bi-directional firing during navigation toward the aperture zone (Assembly #2) and reward zone (Assembly #5) (Figure S9D), or showed goal approaching-related firing increases in both the poke zone (Assembly #3) and the reward zone (Assembly #6) (Figure S9E). The above results demonstrated that the self-information coding concept was useful to discover a variety of prefrontal cell assemblies engaged in distinct stages of five-choice, attentive operant-conditioning task.

### Critical variability-distribution boundaries for constructing efficient assembly code

The ability to uncover a variety of cell assemblies from the multiple brain regions and under multiple conditions and tasks suggest that neural self-information coding represents a general process. The next critical question is whether there is a threshold or boundary in the IS variability distribution that can efficiently signal the shift from the high-probability ground state into the low-probability surprisal states that would give rise to robust assembly-level neural coding. We approached this question by systematically shifting the positive-and negative-surprisal thresholds from 5% to 45% of the skewed ISI gamma distribution tails (Figure 7A). We calculated the cell-assembly coherence index by systematically comparing each assembly’s temporal dynamics obtained from a given threshold with those obtained from all other thresholds. We found that the sliding of the surprisal thresholds or boundaries between 5-15% (ISI variability falling into the low-probability distribution tails) had little effect in terms of improving cell-assembly coherences, whereas assembly-pattern coherence fell apart as the surprisal thresholds shifted from 15% to 30% probability-distribution (Figure 7B). This steep transition around 20% distribution skewed tail was consistently observed all 15 cell assemblies obtained from different brain regions, mental states and behavioral tasks (Figure S10). This strongly suggests that there is a conserved and critical boundary transition for constructing efficient self-information neural codes.

**Figure 7.**
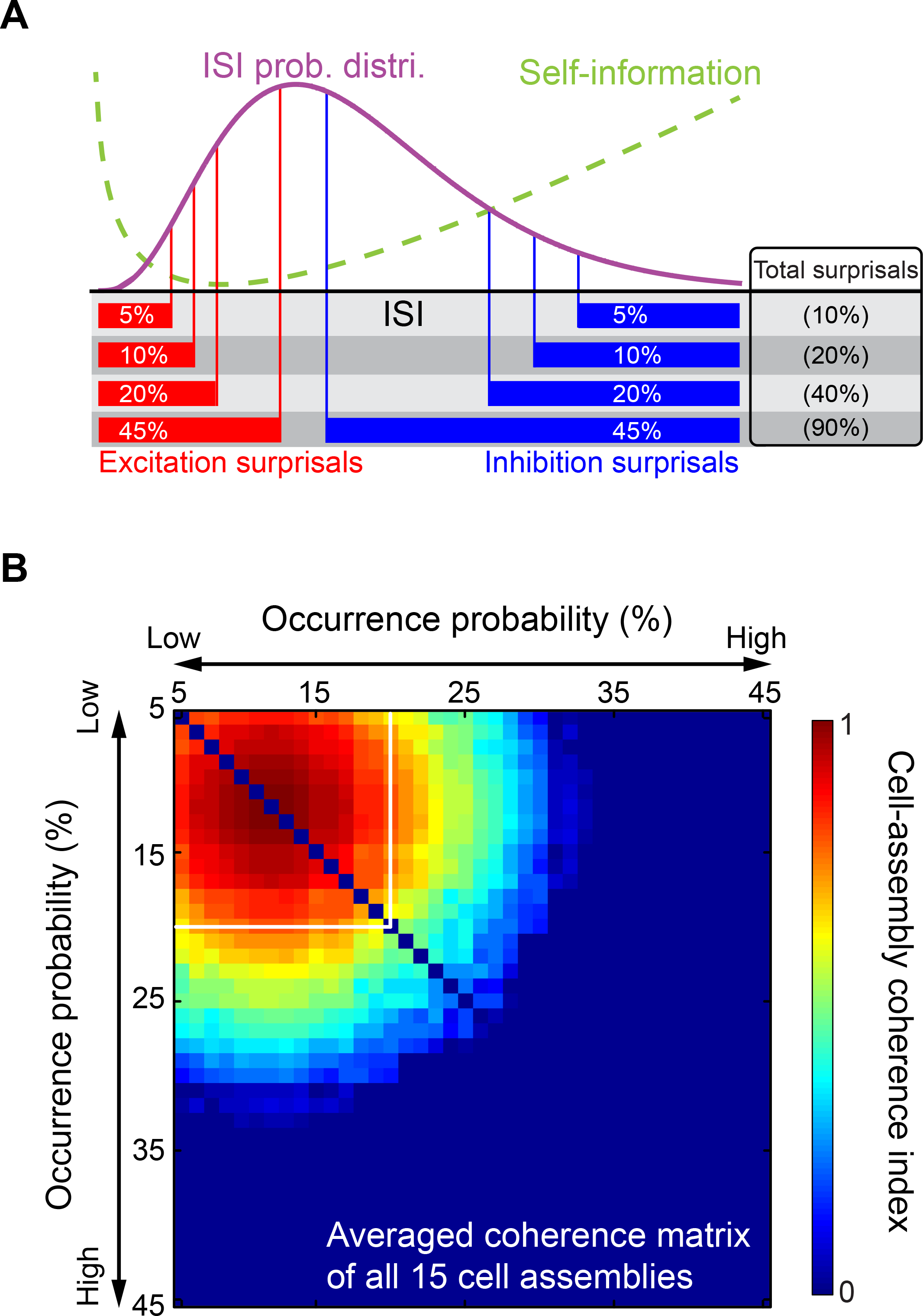
Surprisal boundaries for giving rise to robust real-time cell-assembly codes. (A) Illustration of applying the sliding-window technique to shift the positive-and negative-surprisal thresholds from 5% (low occurrence probability) to 45% (high occurrence probability) of the skewed ISI gamma distribution tails. (B) Shown is the averaged cell-assembly coherence matrix of all 15 cell assemblies. Colors in the matrixes denote corresponding cell-assembly coherence indexes. Robust cell-assembly patterns were observed with an occurrence probability of less than 20% (as denoted by white square frames in cell-assembly coherence matrixes).

### Composition of self-information codes is region-and task-specific

Finally, we explored how dynamic firing changes in putative principal excitatory cells and interneurons would contribute as positive or negative ISI surprisals to construct various cell assemblies (Figure 8A). We first calculated the percentages of positive or negative surprisals out of the total surprisal occurrences in each of the 15 cell-assembly patterns (Figures 8B-E). We then examined how many of these positive or negative surprisals were contributed by the putative excitatory neurons and interneurons. We noted that the majority of cell-assembly members identified in the present study were putative excitatory cells (70.73% in ACC cell assemblies, 78.8% of the total number of CA1 cell assemblies’ members, and 80% in PRL cell assemblies), while the rest were fast-spiking interneurons and unclassified neural types (e.g. units with low firing rates and narrow spike waveforms). As expected, both excitatory neurons and fast-spiking interneurons can contribute to positive and negative surprisals by increasing or decreasing their ISI time durations. However, we found that the percentages of positive and negative surprisals varied dramatically from one assembly pattern to another depending on the type(s) of the nature of mental states or tasks. For example, for coding sleep oscillation states such as theta cycles or ripples, spatial navigation, or processing distinct fearful experiences, these cortical and hippocampal assemblies consisted of overwhelmingly positive surprisals (70~96% of total surprisals) (Figure 8B-D). While excitatory cells contribute overwhelmingly to positive surprisals (due to their large proportion in terms of the total percentage of cell numbers), we noted that fast-spiking interneuron members can be dominant in producing both the positive and negative surprisals in some cases (e.g. CA1 assemblies during sleep).

**Figure 8.**
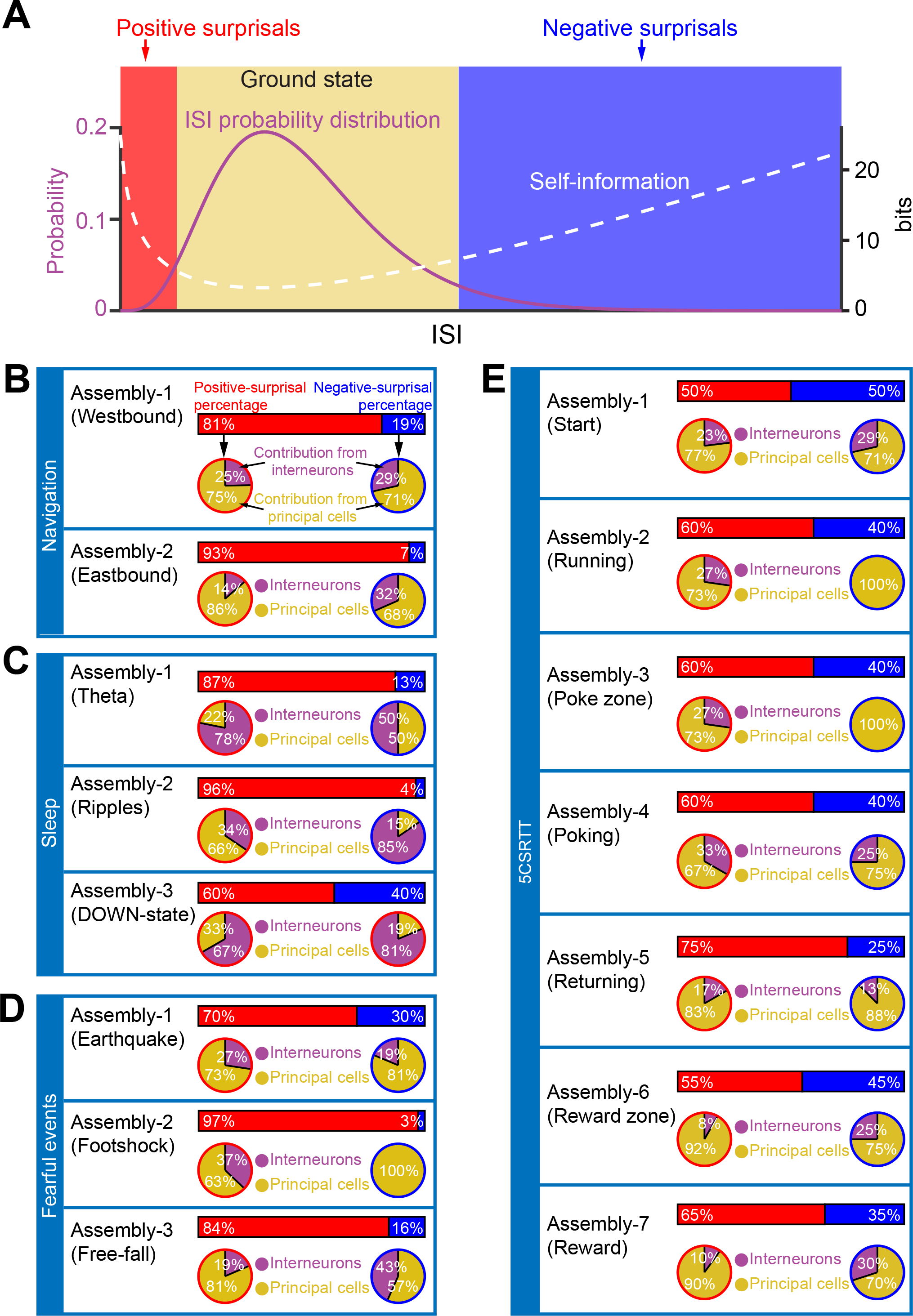
Cell-assembly codes were made of different compositions of positive- and negative-surprisals generated by excitatory cells and fast-spiking interneurons. Based on the variability-surprisal distribution, spike activities of cell-assembly can lead to positive-surprisals or negative-surprisals. (B-E) Percentages of positive- (red bars) or negative-surprisals (blue bars) used to generate distinct cell-assembly codes. The pie charters showed the percentages of positive surprisals (in red bars) were generated from excitatory neurons (yellow) or interneurons (purple). Similarly, the percentages of negative-surprisals (in blue bars) were generated from excitatory neurons (yellow) or interneurons (purple).

Most interestingly, various neural coding constructed by PRL cell-assembly patterns (Figure 8E, Assembly-1, 2, 3, 4, 6, and 7) during a five-choice visual discrimination operant-conditioning task as well as the CA1 Assembly-3 pattern encoding the downstate of sleep cycle (Figure 8C) were constructed by a large proportion of negative surprisals (40~50% of total surprisals). Therefore, the underlying compositions (i.e. positive and negative surprisals, as well as the percentages of excitatory neurons vs. interneurons) were specific to neural circuits, cognitive states, and behavioral tasks.

## Discussion

Here, we tested a novel hypothesis that neuronal variability operates as the selfinformation generator and messenger to convey a variable amount of information in the form of ISI variability surprisal. Coordination of these surprisal ISIs in space (across a population of neurons) and time can intrinsically give rise to robust real-time cell-assembly code (26). This new concept can provide a new conceptual framework to explain how information is robustly signaled in real time by spike trains in the face of enormous neuronal variability. This adds to emerging view in the literature that neuronal variability is not simply synaptic and systems noise (2–5, 8–18, 24, 55–57).

We approached the question of whether firing variability reflects information coding or noise by systematically determining the difference or similarity in neuronal variability across the subcortical and cortical structures. This included the basolateral amygdala, hippocampal CA1, striatum, and three cortical regions such as the RSC, PRL, and ACC. In all cases, we found that variability remained at the similar level, consistent with the prediction that neuronal variability does not reflect the system noise which is expected to become larger as noise would accumulate over each relay. Our present work has also advanced the efforts in extracting the covert structure within the apparent noise in spike trains (9, 24, 55, 56).

One interesting aspect of this self-information coding concept is that the neural code is intrinsic to the neurons themselves (26), with no need for outside observers to set any reference point as typically used in the rate-code, population-code and temporal-code models. A variety of cell assemblies can indeed be identified in an unbiased manner - namely, without traditional reliance on averaging spike trains over external reference points, such as the stimulus delivery time, spatial location, local field potential or specific actions of cognitive behaviors recorded during experiments. Such information or steps were only used after cell-assembly patterns were identified by the VCAD method, as a way to interpret and/or define the potential functions or contribution of each assembly patterns. For analyzing the datasets, we further enhanced the unbiased decoding method by developing a step by which independent-component signal sources was automatically determined using the minimal CV values in each dataset.

To test whether the self-information coding represents a general principle, we applied the VCAD strategy to spike datasets obtained from different brain regions under a set of representative cognitions reflecting the external experiences (encountering earthquake, elevator-drop, foot-shock, navigation and various actions during five-choice visual discrimination tasks) and internal states (sleep oscillation cycles). Selection of these mental states and cognitive tasks also allowed us to examine whether the self-information concept can underlie the encoding of both categorical variables (i.e. distinct fearful stimuli, or nose-poking and obtaining food pellets during the operant-conditioning tasks) and continuous variables (i.e. spatial navigation or sleep). Altogether, we have successfully uncovered 15 different cell assemblies covering a wide range of cognitive processing from the PRL, ACC, and hippocampal CA1.

As the first test-case for the self-information coding concept, we examined whether the VCAD method can be used to uncover cell assembly patterns encoding categorical variables such as earthquake, elevator-drop or mild electric foot-shock experiences. We selected the ACC due to its crucial role in processing emotions and fear behaviors (34,36, 58–60). By extracting self-information coding patterns from firing variability, we identified three distinct cell assemblies that were time-locked to these three fearful events. The contribution of the top ranking members to each cell assembly was quantitatively determined by shuffling experiments (Figure S4A), a feat that was not achievable for principal component analysis (PCA) and multiple-discriminant analysis (MDA) (25). This quantitative assessment of membership information is highly valuable to better understand dynamics and contribution of individual neurons to the overall assembly pattern. Overall, we found that many ACC cells exhibited specific responses to a specific fearful experience (i.e. earthquake vs. foot-shock), whereas some cells responded in a combinatorial manner, such as coding for both earthquake and foot-shock or earthquake and free-fall, etc. Moreover, some ACC cells participated in all cell assemblies, reflecting by their responsiveness to all three fearful events. Such specific-to-general combinatorial cell assemblies in the ACC is highly consistent with the recent finding that the brain is organized by power-of-two permutation-based logic (61, 62).

The inclusion of hippocampal place cells was designed as a gold-standard test-case. Indeed, we showed that by assessing self-information patterns based on spike variability and then simply scanning through the recorded sessions, we uncovered CA1 assembly patterns that were time-locked to either the westbound or eastbound journeys. We showed that each assembly contained classic place cells with a single salient place field (23, 28, 30, 31, 40, 49, 51, 52), but also cells with multiple place fields (63, 64). Unexpectedly, we also found that each assembly consisted of non-place cells such as cells responding preferentially to the start and finish state. Most interestingly, many of these start/finish cells were specific to either the westbound or eastbound journey, suggesting that their firing changes were not simply reflecting generic motion transition. Such properties are consistent with the reports that CA cells may also encode goal-related activity (65–67). Although the current study did not manipulate rewards (no pellets was represented during recording sessions, but only during training trials), the future investigation can be performed to examine such properties. Another unique aspect in uncovering of a variety of CA1 cells associated with spatial navigation behavior is that it provides an unbiased means to potentially investigate a long-held, unresolved question as to how researchers should define place cell assembly and what size a cell assembly might be (28). The top-ranking membership, based on the highest contribution weights derived from VCAD analysis, listed in demixing matrix **W**, can readily enable researchers to identify the ranking numbers as we have illustrated in each plot of Figure 5 and Figure S8.

The third test-case for the validity of self-information coding concept is to uncover cell assemblies that encode internal continuous variables, namely, sleep. We found the existence of three assembly patterns in the CA1 that were time-locked with three sleep oscillation cycles (theta, ripples, and downstate). Traditional classification based spike-phase coupling (theta coupled cells vs non-coupled cells), would predict that all theta-coupled cells may belong to the same cell assembly. Such a logic may also be extended to ripple-coupled cells vs. non-ripple coupled cells. While many theta-coupled or ripple-coupled cells were found in the CA1 datasets as nicely demonstrated in the literature (38–41, 45–48, 50, 52, 68, 69), surprisingly, many of them did not belong to this sleep theta cell assembly or the ripple cell-assembly. Their distinct memberships were further supported by their distinct correlations (Figure S6). Given the reported complexity and diversity of interneurons (45–47) and even pyramidal cell subtypes (70, 71), a future investigation with Cre-lox-mediated neurogenetics (72) and optogenetics (73–75) will be necessary to clearly define the identity of these putative pyramidal and interneuron subtypes.

To further illustrate the conceptual generality and practical value of self-information coding, we investigated neural coding underlying the five-choice visual discrimination operant-conditioning task as the final test-case. This is one of the most classic operant behavioral tasks (76), but rarely studied at neural coding level. This task requires a set of complex cognitive functions, ranging from attentive and preparative behavior, cued-induced precise actions at the right time and right location (76–79). We discovered that the PrL cortex produced seven distinct cell assemblies during the performance of this task. These cell assembly patterns were time-locked to the onset of cues, and cue-induced behavioral actions, with sequential activation of each cell-assembly pattern according to each stage of five-choice visual discrimination operant-conditioning actions. Within the first cell assembly, there are multiple subtypes of cells that exhibited different time courses of responses. One type reacted ramping activity prior to the onset of the cue (light on in the stimulus aperture) (Figure 6B, the bottom PTSH subpanel). This type of PRL cells is consistent with a previous study reporting that activity of some ACC cell corresponded to the level of preparatory (pre-cue) attention in three-choice serial reaction time task in rats (78). Due to the space constraint and the focus of the current study, we limited our analysis to the general usefulness of self-information coding to uncover overall cell assemblies, the detailed analysis of various distinct subtypes of excitatory and interneurons will be highly interesting, especially in light of a recent finding that prefrontal parvalbumin (PV) neurons in control of attention (77). Moreover, also in the ACC using a simpler linear track food foraging task, Kvitsiani et al. demonstrated that the perisomatically targeting PV and the dendritically targeting somatostatin (SOM) neurons had dissociable inhibitory effects (79). A subtype of SOM neurons selectively responded at reward approach, whereas PV neurons responded at reward leaving and encoded preceding stay duration. A more detailed data analysis and selective manipulations, dedicated to the five-choice attentive operant conditioning, will be necessary for future studies to address these important issues.

In the present study, the self-information-based decoding approach revealed that PRL assemblies contained not only a variety of cells processing specific actions and/or
stages of the operant conditioning performances, but also recruited large proportions of negative surprisals to process various information including both categorical-variable patterns (“start,” “poking” and “reward”) and continuous-variable patterns (“running,” “poke zone,” “returning” and “reward zone”) (Figure 8E). This may provide a new insight into a broader picture regarding the cell-assembly nature of the inhibitory control performed by the prefrontal cortex (80, 81). Such a composition as seen in the PRL cortical assemblies in this attentional operant-conditioning tasks seemed to be different from the encoding of fearful experiences in the ACC (Figure 8D). This is consistent with the studies showing that fearful stimuli often increased firing or bursting in the prefrontal neurons (58, 60, 82). Such a difference in compositions in surprisal types and cell types strongly suggest that construction of self-information code is dependent on the tasks and brain regions. The difference in the positive vs. negative surprisal make-up is further illustrated in the cell assemblies related to sleep oscillations, which engaged much more interneurons (Figure 8C). Taken together, these observations provide the first comparative pictures regarding the underlying compositions of cell assemblies, open the door to detailed and systematic investigations into the nature of cell-assembly coding specific to neural circuits, cognitive states, and behavioral tasks.

Finally, as a critical step to further understand how self-information code rises dynamically from neuronal variability, we investigated whether there is a critical boundary in transitioning ISI variability from high probability, low information state to low probability, high information state. Systematic analyses of all 15 cell assemblies using the sliding window technique revealed a steep transition in using ISI surprisals, around 20% of the skewed ISI gamma distribution tails, to generate efficient real-time cell-assembly codes. In another word, 20% of ISIs represented as low-probability surprisals for the construction of various population-level information patterns, whereas the majority (80%) of remaining ISIs operate as the on-going low-information ground states. This 80/20 statistical rule is surprisingly conformed to the Pareto Principle (83), also known as the principle of factor sparsity, which states that, for many events, roughly 80% of the effects or consequences (i.e. information) stem from 20% of the causes (i.e. signs or entropy). This principle has been widely executed in a variety of systems, ranging from economics to software engineering and communications in which optimal data streams are implemented by the minimization of the numbers of signs while maximizing the transmitted information (83–86). It is noteworthy that while the ground state corresponds to the most probable ISI which carry less self-information but may still consume a lot of energy, these ground-state ISIs can play an extremely important role in enabling both the rapid responses to changes and ternary coding structure once combined with positive and negative surprisals (26). This ternary coding dynamics can lead to enormous information capacity and greater operational flexibility for neural clique assemblies which are reportedly organized by power-of-two-based permutation logic across various brain regions (61, 62).

In summary, we show that neuronal variability operates as a self-information generator and messenger to produce real-time ISI variability surprisals that can be used unbiasedly to uncover various cell assemblies corresponding to both categorical and continuous variables. These self-information-based neural codes are uniquely intrinsic to the neurons themselves, with no need for outside observers to set any reference point to manually mark external or internal inputs. Analysis of 15 distinct cell assemblies suggests the existence of conserved critical boundaries, seemly conforming to the Pareto principle, in utilizing ~20% of ISIs as statistical variability surprisals from which robust real-time cell-assembly patterns arise efficiently. The unique ability to identify numerous cell assemblies based on the unified self-information principle provided us with a rare and unbiased assessment for the compositions of cell-assembly codes.

## Materials and Methods

### Ethics Statement

All animal work described in the study was carried out in accordance with the guidelines laid down by the National Institutes of Health in the United States, regarding the care and use of animals for experimental procedures, and was approved by the Institutional Animal Care and Use Committee of Augusta University (Approval AUP number: BR07-11-001).

### I*n vivo* recording in mice and data processing

Tetrodes and headstages were constructed using the procedures as we have previously described (62, 87). The CA1 units were recorded male wild-type B6BCA/J mouse using the adjustable 96-channel recording array (with 48-channels bilaterally), as previously described (46), whereas 128-channel tetrode microdrives were used to record from the BLA, dorsal striatum, ACC, RSC, PrL bilaterally with 64 channels per hemisphere. For facilitating the identification of electrode array position, the electrode tips were dipped in fluorescent Neuro-Dil (Neuro-DiI, #60016, Red oily solid color, from Biotium, Inc.) which then can reveal the electrode track.

Neuronal activities were recorded by MAP system (multi-channel acquisition processor system, Plexon Inc., Dallas, TX) in the manner as previously described (46). Extracellular action potentials and local field potentials data were recorded simultaneously and digitized at 40 kHz and 1 kHz respectively. The artifact waveforms were removed and the spike waveform minima were aligned using the Offline Sorter 2.0 software (Plexon Inc., Dallas, TX), which resulted in more tightly clustered waveforms in principal component space. Spike-sortings were done with the MClust 3.3 program with an autoclustering method (KlustaKwik 1.5). Only units with clear boundaries and less than 0.5% of spike intervals within a 1 ms refractory period were selected. The stability of the in vivo recordings was judged by waveforms at the beginning, during, and after the experiments. Well-separated neurons were assessed by “Isolation Distance” and “Lratio” (88). Neurons whose “Isolation Distance” >15 and “L-ratio” <0.7 were selected for the present analyses.

### Variability-based cell assembly discovery (VCAD) method

Three well-defined statistics were adopted to quantitatively describe the variability of the neuron’s ISI - namely, the coefficient of variation (*CV*), skewness and kurtosis (see Supporting Information). The VCAD involves three distinct steps (also see Supporting Information): The first step was to use gamma distribution model to describe the variability of the neuron’s ISI (33). The second step was to use the independent-component analysis (ICA) method to decode the ensemble patterns from the population surprisal code by searching joint variability-surprisals. The third step was to identify corresponding cell assemblies by examining top large-weight neurons, which can be directly identified by their weights in demixing matrix **W** of ICA analysis.

### Fearful events and corresponding data analysis

Mice were subjected to three fearful episodic events, earthquakes, foot-shocks, and free-fall drops (see Supporting Information for detailed description). These episodic stimuli are fearful as evidenced from physiological indications including a rapid increase in heart rates as well as reduced heart rate variability (37, 89). To maintain the consistency of stimulation timing (minimizing the possible prediction of upcoming stimuli), the stimuli were triggered by a computer and delivered at randomized intervals within 1-3 minutes. After the completion of all fearful event sessions, the mouse was placed back into the home cages.

### Sleep and corresponding data analysis

Data was recorded when the mouse was sleeping in the home cage. The recorded local field potentials (LFP) were first processed by the FPAlign (a utility program provided by Plexon Inc.) to correct the filter induced phase delays. LFP channels recorded in CA1 pyramidal cell layer were selected by judging the maximum yield of ripple during animal slow-wave sleep (SWS), and comparing the coherence in the theta-frequency and gamma-frequency band (90). Further analyses were carried out with these LFP data offline by custom-written MATLAB (Mathworks) programs. The local field potential was band passed at Delta (2-4 Hz), Theta (4-10 Hz), low Gamma (25-90 Hz), fast Gamma (90-130 Hz) and hippocampal ripples (130-200 Hz) by hamming window based FIR filters with order of 30 (see Supporting Information for detailed description).

### Linear track and corresponding data analysis

A linear track experiment used a 100cm×10cm closed runway. The exploration duration was 20 minutes. Real-time positions of the mouse were tracked by CinePlex Behavioral Research Systems (Plexon Inc., Dallas, TX) automatically. Place fields were defined as a set of ≥5 contiguous pixels with a firing rate above two standard deviations of the mean firing rate. All spatial firing plots were smoothed with 5 × 1 Gaussian smoothing filter using the Matlab software (see Supporting Information for detailed description).

### Five-choice visual discrimination operant-conditioning task and corresponding data analysis

The animal was introduced to a custom-designed test chamber equipped with five response apertures that can illuminate, and a food magazine to deliver reward. A trial was initiated by the mouse entering the food magazine. A brief light-stimulus was then presented in one of five possible apertures after a 5-s inter-trial interval (ITI) had elapsed.

The mouse was required to scan the five apertures for the appearance of the light-stimulus and to respond in the “correct” aperture with a nose-poke response in order to earn a single food pellet delivered in the food magazine. If the mouse responded before the stimulus (“premature response”) or in an adjacent, incorrect aperture (“incorrect response”), a 5-s time-out (TO) period was introduced by extinguishing the house light and not providing a food reward. Failure to respond within the limited hold (LH) period resulted in an ‘omission” and a subsequent 5-s TO period. After collecting the reward or - on punished trials - at the end of the TO period, a head entry in the food magazine would start a new trial. The mouse was pre-trained to achieve ≥ 50 correct trials within 30 minutes with >80% accuracy and <20% omissions before recording from prefrontal cortex began. Animal’s movements were recorded by CinePlex Behavioral Research Systems (Plexon Inc., Dallas, TX), and the animal’s locations and head directions were tracked manually afterward. Detailed protocol was described in Supporting Information.

## Author Contributions

J.Z.T. conceived and designed the project with M.L., J.Z.T. designed the experiments with M.L., K.X., H.K., G.E.F., and J.L., The research was performed as follows: K.X. recorded from the ACC and PRL datasets; G.E.F. from the RSC; Jun Liu from the BLA; D.W. from the striatum, H.K. for the CA1. M.L., F.Z., H.K., Z.S., L.C. and Y.M. for data analyses together with J.Z.T., J.Z.T. and M.L. wrote the paper with input from all other.

## Acknowledgments

This work is supported by an NIH grant (R01NS079774), GRA equipment grant to J.Z.T., Shanghai Youth Science and Technology Sail Project (16YF1415200) to Z.S., M.L. and the Brain Decoding Center grant Yunnan Science Commission (2014DG002) to F.Z. and J.Z.T. We thank Fengying Huang for maintaining the mouse colony and training mice for five-choice operant-conditioning tasks, and Sandra E. Jackson for proofreading this manuscript.

## Supplementary information

### SI Materials and Methods

#### In *vivo* recording in mice and data processing

Tetrodes and headstages were constructed using the procedures as we have previously described (1, 2). To construct tetrodes, a folded piece consisting of four wires (90% platinum, 10% iridium, 13 μm, California Fine Wire Company, Grover Beach, CA, USA) was twisted together using a manual turning device and soldered with a low-intensity heat source (variable temperature heat gun 8977020, Milwaukee, Brookfield, WI, USA) for 6 s. The impedances of the tetrodes were measured with an electrode impedance tester (Model IMP-1, Bak Electronics, Umatilla, FL, USA) to detect any faulty connections, and our tetrodes were typically between 0.7 MΩ and 1 MΩ. The insulation was removed by moving the tips of the free ends of the tetrodes over an open flame for approximately 3 s. The tetrodes were then placed into appropriate polyimide tubes. The recording ends of the tetrodes were cut differentially (Vannas spring scissors -3 mm cutting edge, Fine Science Tools, Foster City, CA, USA) according to the different depths of the recording sites. This ensures that only tetrodes, but not the surrounding polyimide tubes, were inserted into the brain tissue, thereby minimizing the tissue damage.

We employed adjustable 128-channel tetrode microdrives to target the basolateral amygdala (BLA; n = 8 WT mice), anterior cingulate cortex (ACC; n = 7 WT mice), the retrosplenial cortex (RSC; n = 20 WT mice), dorsal striatum (STR; n = 7 WT mice), and prelimbic region (PrL; n = 4 WT mice) bilaterally with 64 channels per hemisphere. Whereas the CA1 units were recorded from nine WT mice using the adjustable 96-channel recording array (with 48-channels bilaterally). Stereotaxic coordinates were as follows: for BLA, 1.7 mm posterior to bregma, 3.5 mm lateral, -4.0 mm ventral to the brain surface; for ACC, +0.5 mm AP, 0.5 mm ML, -1.75 mm DV; for CA1, 2.0 mm lateral to the bregma and 2.3 posteriors to the bregma; for RSC, -2.5 mm AP, 0.5 mm ML, -0.8 mm DV; for STR, +0.62 mm AP, 1.35 mm ML, -2.0 mm DV; and for PrL, +1.70 mm AP, ± 0.5 mm ML (3).

Male wild-type mice (6-8 months old) were moved from home cages housed in the LAS facility to the holding area next to the chronic recording rooms in the laboratory and stayed in a large plastic bucket (20 inches in diameter and 16 inches in height - per mouse, Walmart) with access to water and food for a week prior to surgery. During this period, the animals were also handled daily to minimize the potential stress from human interaction. The animal was given an intraperitoneal injection of 60 mg/kg ketamine ( Bedford Laboratories, Bedford, OH, USA) and 4 mg/kg Domitor (Pfizer, New York, NY, USA) prior to the surgery. Animal’s head was secured in a stereotaxic apparatus, and an ocular lubricant was used to cover the eyes. The hair above the surgery sites was removed, and Betadine solution was applied to the surface of the scalp. An incision was then made along the midline of the skull. Hydrogen peroxide (3% solution, Fisher Scientific) was placed onto the surface of the skull so that bregma could be visualized. The correct positions for implantation were then measured and marked. For fixing the microdrive headstage, four holes for screws (B002SG89S4, Amazon, Seattle, WA, USA) were drilled on the opposing side of the skull and, subsequently, the screws were placed in these holes with reference wires being secured to two of the head screws. Craniotomies for the tetrode arrays were then drilled, and the dura mater was carefully removed. After the electrodes were inserted and tetrodes were secured to the fiberglass base, the reference wires from the connector-pin arrays were soldered such that there would be a continuous circuit between the ground wires from the head screws and those from the connector-pin arrays. Finally, the connector-pin arrays were coated with epoxy. Aluminum foil was used to surround the entire headstage to aid in protection and to reduce noise during recordings. The animals were then awoken with an injection of 2.5 mg/kg Antisedan. The animals were allowed to recover post-surgery for at least 3-5 days before recording. Then, the electrode bundles targeting the BLA, STR and hippocampal CA1 region were slowly advanced over several days in small daily increments. For the cortical sites, tetrodes were advanced usually only once or twice in a small increment. At the end of the experiments, the mice were anesthetized and a small amount of current was applied to the recording electrodes in order to mark the positions of the stereotrode bundles. The actual electrode positions were confirmed by histological Nissl staining using 1% cresyl echt violet. For facilitating the identification of electrode array position, the electrode tips were dipped in fluorescent Neuro-Dil (Neuro-DiI, #60016, Red oily solid color, from Biotium, Inc.) which then can reveal the electrode track.

Neuronal activities were recorded by MAP system (multi-channel acquisition processor system, Plexon Inc., Dallas, TX) in the manner as previously described (4). Extracellular action potentials and local field potentials data were recorded simultaneously and digitized at 40 kHz and 1 kHz respectively. The artifact waveforms were removed and the spike waveform minima were aligned using the Offline Sorter 2.0 software (Plexon Inc., Dallas, TX), which resulted in more tightly clustered waveforms in principal component space. Spike-sortings were done with the MClust 3.3 program with an auto-clustering method (KlustaKwik 1.5). Only units with clear boundaries and less than 0.5% of spike intervals within a 1 ms refractory period were selected. The stability of the in vivo recordings was judged by waveforms at the beginning, during, and after the experiments. Well-separated neurons were assessed by “Isolation Distance” and “Lratio” (5). Neurons whose “Isolation Distance” <15 and “L-ratio” >0.7 were excluded in the present analyses.

#### Analysis of ISI’s variability

Three well-defined statistics were adopted to quantitatively describe the variability of the neuron’s ISI - namely, the coefficient of variation (*CV*), skewness and kurtosis. The definitions of these three variability measurements were as follow:

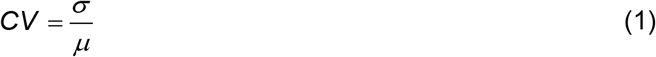

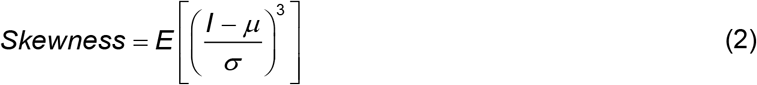

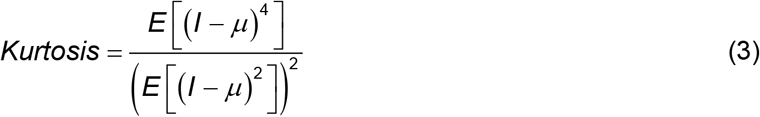

where *I* = (*i*_1_,*i*_2_,…,*i*_*n*_) denotes ISI of a neuron, *σ* is the standard deviation of *I*, and *μ* is the mean of *I*. As shown in Figure 1B, BLA dataset has 496 putative principal cells from 8 mice, CA1: 495 putative principal cells from 9 mice, STR: 295 putative principal cells from 7 mice, RSC: 504 putative principal cells from 5 mice, PrL: 85 putative principal cells from 4 mice, ACC: 195 putative principal cells from 6 mice.

We also conducted the same analysis to two datasets recorded from BLA and hippocampal CA1 under two distinct states - namely awake state and anesthesia. Recordings were first carried out when animals were awake and immobile in their home cages. Minimum 40-minute neural activities were recorded for awake state analysis. To produce ketamine-induced anesthesia, the animals were injected with a 60 mg/kg ketamine and 0.5 mg/kg Domitor cocktail mixture via i.p., the animals lost the righting reflex in a few minutes. Neural spike activities were recorded for 50 minutes under anesthetized state. 40-minute neural spike data recorded from the fully anesthetized state starting from 10 minutes after the Ketamine/Domitor injections were selected for the present analysis. For neuronal variability analyses of awake state vs. anesthesia (Figure 1C), the CA1 dataset contains 59 putative principal cells from 3 mice, and the BLA dataset has 92 putative principal cells from 2 mice.

#### Variability-surprisal based cell assembly discovery (VCAD) framework

##### 1) Assessing ISI’s variability.

In this step, a gamma distribution model was applied to describe the variability of the neuron’s ISI (6). A gamma distribution model is defined by a shape parameter *k* > 0 and a scale parameter *θ* > 0, and the gamma probability density function for spike trains is defined by

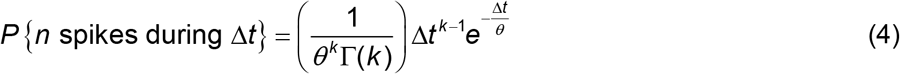

where Γ(*k*) stands for the gamma function.

We fitted the gamma distribution model for the ISIs of each neuron. Shape parameter *k* and scale parameter *θ* were estimated by the maximum likelihood estimates (MLE) method, which selected values of these two parameters that produced a gamma distribution for the ISIs with minimal errors. To avoid an under-sampled error due to too few data points, units with <250 ISIs were excluded in the present analysis.

For each cell, two boundaries of the ground state were assigned base on the fitted gamma distribution. Using these two thresholds, the neural spike trains were converted to variability surprisals - that is, positive surprisals, negative surprisals and ground state. Specifically, we first binned the neural spike trains (bin size = 200ms for all analyses), and the ISI of the bin *ISI*_*bin*_ was set as follows:

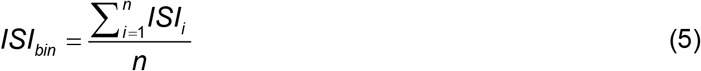

where *n* is the total number of ISIs which occurred in the bin (including partly-occurred ISIs), *ISI*_*i*_ is the time duration of the *i*th ISI in the bin. The bins whose ISIs are less than the lower boundary of ground state were assigned as positive surprisals (red region in the middle subpanel of step 1 in Figure 2A, which means shorter ISIs and higher instant firing rates), bins whose ISIs are larger than the higher boundary of ground state were assigned as negative surprisals (blue region in the middle subpanel of step 1 in Figure 2A, corresponding to longer ISIs and lower instant firing rate), and all other bins were assigned as ground state. In this way, the neuron’s spike train was converted into a surprisal code. In the present analysis, the boundaries of ground state were set as 10% and 90% based on the fitted gamma CDF curves - that is, the ISIs whose gamma CDF is less than 10% or higher than 90% were assigned as positive or negative surprisals, respectively.

##### 2) Uncovering joint surprisal patterns

In this step, the blind source separation (BSS) method was applied to decode the ensemble patterns from the population surprisal code by searching joint variability-surprisals. Independent Component Analysis (ICA, a BSS technique) was adopted to unbiasedly uncover distinct ensemble patterns; that is, reducing high-dimensional data (population surprisal code) to a set of independent information sources corresponding to different brain states.

ICA is a statistical technique that aims at finding a separating or demixing matrix **W** for generating linear projections of the input data that maximize their mutual independence. The ICA model assumes that the recorded neuronal population spike activities (represented by **SA**(*t*) = [*sa*_1_(*t*),*sa*_2_(*t*),…,*sa*_*n*_(*t*)]) contain mixtures of the *n* underlying information sources (represented by **IS**(*t*) = [*is*_1_(*t*),*is*_2_(*t*),…,*is*_*n*_(*t*)]),i.e.,

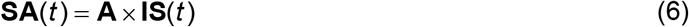

where the square matrix **A** contains the mixture coefficients *a*_*ij*_. The aim of ICA is to find a demixing matrix **W** that is an approximation of the inverse of the original mixing matrix **A** whose output

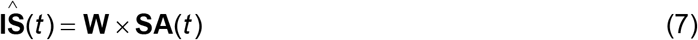

is an estimate of **IS**(*t*) containing the underlying information sources (ensemble patterns of corresponding cell assemblies). To uncover the independent information sources, **W** must maximize the non-Gaussianity of each source. In practice, iterative methods are used to maximize or minimize a given cost function that measures non-Gaussianity. ICA analyses were performed by FastICA Matlab package that implemented the fast fixed-point algorithm (7).

The major challenge of BSS analysis of neural population activities is how to select the optimum number of information sources that provide the best representation of distinct brain states. The nature of the reduced dimension *p* means that *p* information sources are extracted from neural population activities. Many subjective methods can be derived but ultimately the relevance of each ensemble pattern depends on the nature of data being analyzed.

Here we introduced a criterion for selecting the best relevant ensemble patterns. This criterion was based on the hypothesis that cell-assembly coding under a given cognitive state should be most efficiently achieved by using a minimal amount of energy consumption while producing a sufficient robustness in the population-encoding of information. That is to say, the assembly code should use the least amount of variability changes of individual cells together with the minimal numbers of such information-coding cells. Thus, for all possible *p*, we calculated the number of the coefficient of variation (*CV*) of the numbers of large-contributed neurons (neurons whose weights in demixing matrix **W** exceeded three times the standard deviation (S.D.) of all weights) over all channels. For example, supposing *p* = 6 which meant there were six ICs, and the numbers of large-contributed neurons in six ICs were{6,8,11,7,7,5}, the mean and S.D. of these numbers were 7.33 and 2.07, therefore the *CV* for *p* = 6 was 0.28 (2.07/7.33). In this way, a vector {*CV*_*i*_} was produced, where *i* covered all possible values of *p*. The dimension with min({*CV*_*i*_}) was unbiasedly selected as the best relevant dimension. An example is shown in Figure S3A, {*CV*_*i*_} were calculated for a set of neural population activity with *p* from 2 to 20, and dimension 3 with min ({*CV*_*i*_}) was chosen (highlighted by the red circle).

##### 3) Identifying corresponding cell assemblies

For each decoded information source (ensemble pattern), the members of the corresponding cell assembly were selected as top large-weight neurons, which can be directly identified by their weights in demixing matrix **W** of ICA analysis. It should be noted that cells’ weights in **W** could be either positive or negative values; here, we used absolute values of cells’ weights for the identification of corresponding cell assemblies.

#### Fearful events and corresponding data analysis

Mice were subjected to three fearful episodic events, earthquakes, foot-shocks, and free-fall drops. For the earthquake-like shake, the mouse was placed in a small chamber (4” × 4” × 6“H circular chamber) fixed on top of a vortex mixer and shaken at 300 rpm for 400ms for six times with 1~3-minute time interval between each shake. For fearful foot-shock, the foot-shock chamber was a square chamber (10” × 10” × 15“H) with a 24-bar shock grid floor. The mouse was placed into the shock chamber for three minutes and received the foot-shock stimulus (a continuous 300-ms foot-shock at 0.75 mA) for a total of six times with inter-trial time interval between 1~3 minutes. For free-fall in the elevator, the animal was placed inside a small box (3” × 3” × 5“H) and dropped from a 13-cm height (a cushion which made from a crumbled table cloth was used to dampen the fall and to stop the bouncing effect). After 1~2 minutes, the elevator was raised gently back to the 13-cm height and dropped again after 1~2 minutes (this process was also repeated six times). These episodic stimuli are fearful as evidenced from physiological indications including a rapid increase in heart rates as well as reduced heart rate variability (8, 9). To maintain the consistency of stimulation timing (minimizing the possible prediction of upcoming stimuli), the stimuli were triggered by a computer and delivered at randomized intervals within 1-3 minutes. After the completion of all fearful event sessions, the mouse was placed back into the home cages.

#### Sleep and corresponding data analysis

Data was recorded when the mouse was sleeping in the home cage. The recorded local field potentials (LFP) were first processed by the FPAlign (a utility program provided by Plexon Inc.) to correct the filter induced phase delays. LFP channels recorded in pyramidal cell layer were selected by judging the maximum yield of ripple during animal slow-wave sleep (SWS), and comparing the coherence in the theta-frequency and gamma-frequency band (10). Further analyses were carried out with these LFP data off-line by custom-written MATLAB (Mathworks) programs. The local field potential was band passed at Delta (2-4 Hz), Theta (4-10 Hz), low Gamma (25-90 Hz), fast Gamma (90-130 Hz) and hippocampal ripples (130-200 Hz) by hamming window based FIR filters with order of 30.

For extraction of ripple events, the root mean square power of the 130-200 Hz filtered LFP was calculated by sliding 10-ms window every one millisecond and averaged across electrodes. A threshold was set to five standard deviations above the mean power to detect ripples. The beginning and the end of oscillatory epochs were marked at points with the power reduced to two standard deviations above background mean. Periods of 20-ms or more were designated ripple episodes. The minimum point during each episode was regarded as the reference to calculate the peri-event histogram for each individual unit. If there are at least 3 bins during -30ms to 30ms (bin size: 10ms) greater than two standard deviations above the mean (the -500ms to -100ms and 100ms to 500ms epochs were used as a baseline), then this unit was considered having significantly elevated firing rate during ripple episodes.

For detection of theta epochs, the theta/delta power ratio was calculated in a 2s window. More than consecutive 15s periods in which the ratio was greater than 4 were identified, then the beginnings and ends of epochs were judged manually. To calculate the phase relationship between unit activity and theta oscillation, the LFP data was first digitally band-bass filtered (4-12 Hz). The troughs for each cycle were identified and were assigned instantaneous phase of 0 radians. The phase value of each spike was obtained by linear interpolation. Then the histogram was calculated with 20-degree bin size, and a Rayleigh’s circular statistics was performed to determine the significance of phase modulation of theta oscillations.

#### Linear track and corresponding data analysis

A linear track experiment used a 100cm×10cm closed runway. The exploration duration was 20 minutes. Real-time positions of the mouse were tracked by CinePlex Behavioral Research Systems (Plexon Inc., Dallas, TX) automatically.

For each recorded unit, a corresponding 100x1 array of time-averaged firing rates was generated by dividing, on a pixel-by-pixel basis (1cm×10cm), the number of spikes by the time in each location. For place cell (including place cells with single or multiple fields) and differential cell analysis, a motion speed threshold was applied to generate the time-averaged firing rates arrays (i.e., the spikes were excluded if collected from an animal moving at speeds of less than 2.5 cm/s). Place fields were defined as a set of ≥5 contiguous pixels with a firing rate above two standard deviations of the mean firing rate. For start/finish cell analysis, no speed limit applied, and significant firings at two ends of the line track were defined as a set of ≥5 contiguous pixels with a firing rate above two standard deviations of the mean firing rate. All spatial firing plots shown in Figure 3 were smoothed with 5×1 Gaussian smoothing filter using the Matlab software.

#### Five-choice visual discrimination operant-conditioning task and corresponding data analysis

The animal was introduced to a custom-designed test chamber equipped with five response apertures that can illuminate, and a food magazine to deliver reward ( Med Associates, Inc., VT. Product numbers: DIG-703B-USB, ENV-203-20, ENV-115C, ENV-312, SG-716B, ENV-307W, and ENV-312-2W). On the front wall of the chamber were five holes equipped with infrared (IR) detectors. IR photo beams were also presented inside the food magazine fitted on the opposite wall, where the house light was located.

The floor of the chamber consisted of stainless steel rods, and there was a removable tray at the bottom. The pellet dispenser was located outside of the box and automatically delivered food pellet to the magazine through a plastic tube. Nose-poking responses were detected by a sensitive infrared detector placed 1/4” in from the front edge of each unit. The reward pellets were delivered automatically when the correct nose-poking responses were detected. All experimental procedures were controlled by Five Choice Serial Reaction Time Task Utility ( Med Associates, Inc., VT).

A trial was initiated by the mouse entering the food magazine. A brief light-stimulus was then presented in one of five possible apertures after a 5-s inter-trial interval (ITI) had elapsed. The mouse was required to scan the five apertures for the appearance of the light-stimulus and to respond in the “correct” aperture with a nose-poke response in order to earn a single food pellet delivered in the food magazine. If the mouse responded before the stimulus (“premature response”) or in an adjacent, incorrect aperture (“incorrect response”), a 5-s time-out (TO) period was introduced by extinguishing the house light and not providing a food reward. Failure to respond within the limited hold (LH) period resulted in an “omission” and a subsequent 5-s TO period. After collecting the reward or - on punished trials - at the end of the TO period, a head entry in the food magazine would start a new trial.

The mouse was pre-trained to achieve ≥ 50 correct trials within 30 minutes with >80% accuracy and <20% omissions. After pre-training, 20-minute neural activity was recorded from prefrontal cortex (specifically, prelimbic cortex) during 5CSRTT. During recording, the correct responses were made within 5.81 ± 0.71 s and reward food pellets were collected within 4.54± 0.29s after correct nose-pokings. Animal’s movements were recorded by CinePlex Behavioral Research Systems (Plexon Inc., Dallas, TX), and the animal’s locations and head directions were tracked manually afterward.

For each neuron, a corresponding 40×25 array of time-averaged firing rates was generated by dividing, on a pixel-by-pixel basis (1cm×1cm), the number of spikes by the time in each location (the spikes were excluded if collected from animal moving at speeds less than 2.5 cm/s). Spatial-specificity firing patterns were defined as a set of ≥9 contiguous pixels with a firing rate above two standard deviations of the mean firing rate. The Spatial-specificity firing maps shown in Figure 6 were smoothed with 6×6 Gaussian smoothing filter using the Matlab software.

We also calculated an orientation-firing polar histogram for each unit. The orientation-firing polar histograms were generated by dividing the number of spikes fired when the mouse faced a particular direction (in bins of 10°) by the total amount of time the mouse spent facing that direction in the task chamber. No smoothing was applied to the resulting circular distribution. Rayleigh’s circular statistics was applied to determine the significance of directional tuning of units.

#### Cell-assembly coherence index

In the present analyses, we set the boundaries of ground state as 10% and 90% of the CDF of the skewed distribution model to uncover cell assemblies, and the decoded cell assemblies verified the effectiveness of these two boundaries. However, can robust cell-assembly decoding results be achieved with other settings of boundaries? To explore these questions, we examined the cell-assembly coherence index between decoded ensemble patterns under different boundaries of the ground state.

Specifically, we systemically varied lower boundaries from 5% to 45% with step by 1%, the corresponding higher boundaries were set as one minus the value of lower boundaries accordingly (from 95% to 55%, as shown in Figure 7A). Taking demixing matrix **W** generated by boundaries of 10% and 90% as a standard decoder, a set of ensemble patterns was then generated with different boundaries for each cell assembly. We then calculated the Pearson correlation coefficients between these ensemble patterns corresponding to same state/stimuli. The Pearson correlation coefficient *ρ*(*EP*_*A*_,*EP*_*B*_) between ensemble pattern *EP*_*A*_ and ensemble pattern *EP*_*B*_ is defined as:

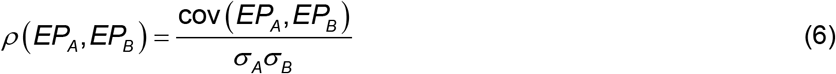

where cov (*EP*_*A*_, *EP*_*B*_) is the covariance of *EP*_*A*_ and *EP*_*B*_, *σ*_*A*_ and *σ*_*B*_ are the standard deviation of *EP*_*A*_ and *EP*_*B*_, respectively.The cell-assembly coherence matrix of each cell assembly is generated by normalizing all corresponding correlation coefficients into range of 0 to 1.

#### Test of statistical significance

In the present study, one-way ANOVA analysis and Tukey post hoc tests were conducted for the comparisons of multiple means. Student’s t-test was used to assess whether two sets of data were significantly different from each other. In all figures that contained statistical significant test results, one asterisk denoted that the ***P*** value is in the range of 0.05-0.01, two asterisks denoted that the ***P*** value is in the range of 0.01-0.001, three asterisks denoted the ***P*** value is less than 0.001. Data were represented as mean ± S.E.M.

**Figure S1.**
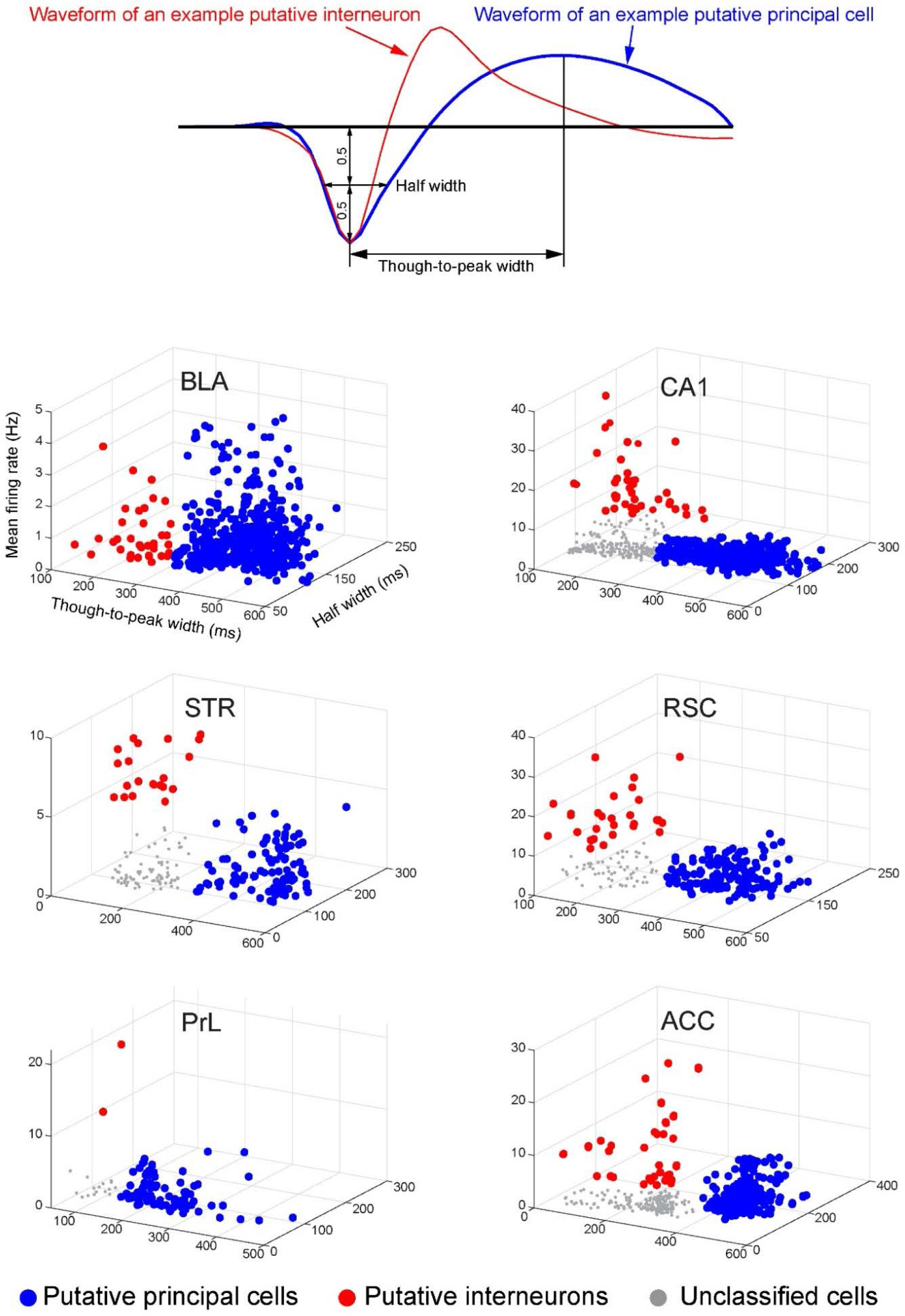
Classification of putative principal cells and interneurons in 6 different brain regions. Isolated units are classified as putative excitatory principal cells or interneurons based on their characteristic firing activity including spike waveforms and firing rates. In general, putative principal cells fire at lower rates and have broader waveforms, whereas, interneurons have higher rates and relatively narrower waveforms. Numbers of putative principal cells: for BLA, 496 principal cells out of 601 recorded units; for CA1, 495 principal cells out of 713 recorded units; for STR, 295 principal cells out of 359 recorded units; for RSC, 504 principal cells out of 651 recorded units; for PrL, 85 principal cells out of 119 recorded units; and for ACC, 195 principal cells out of 397 recorded units.

**Figure S2.**
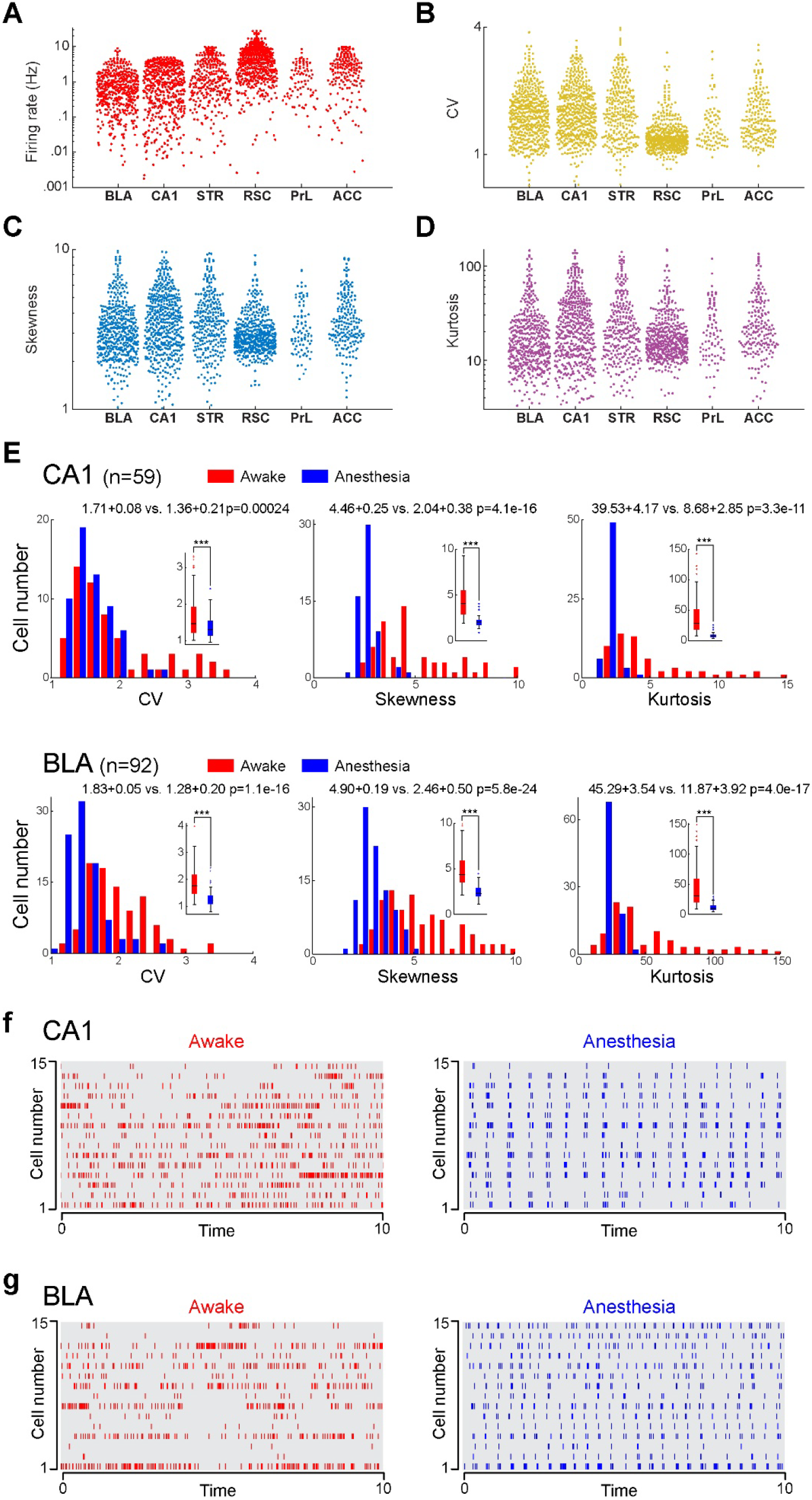
Neural variability analysis across different regions. (A) Distributions of firing rates of principal neurons in six brain regions. (B) Distributions of CV of principal neurons in six brain regions. (C) Distributions of Skewness of principal neurons in six brain regions. (D) Distributions of Kurtosis of principal neurons in six brain regions. All data shown in A-D is in logarithmic coordinates. It should be noted that the CV, Skewness, and Kurtosis in all these six brain regions show Gaussian-like distributions under logarithmic coordinates, indicating that the “long-tailedness” of the ISI distributions also follow skewed statistics. (E) Shown are the histograms of CV, Skewness, and Kurtosis of ISI of putative principal cells in BLA and CA1 under two states - namely, awake state and ketamine-induced anesthesia. Compared with awake state, all these three statistics decrease significantly under anesthesia in both brain regions. (F) Spike activities of 15 CA1 cells during awake state and anesthesia. (G) Spike activities of 15 BLA cells during awake state and anesthesia.

**Figure S3.**
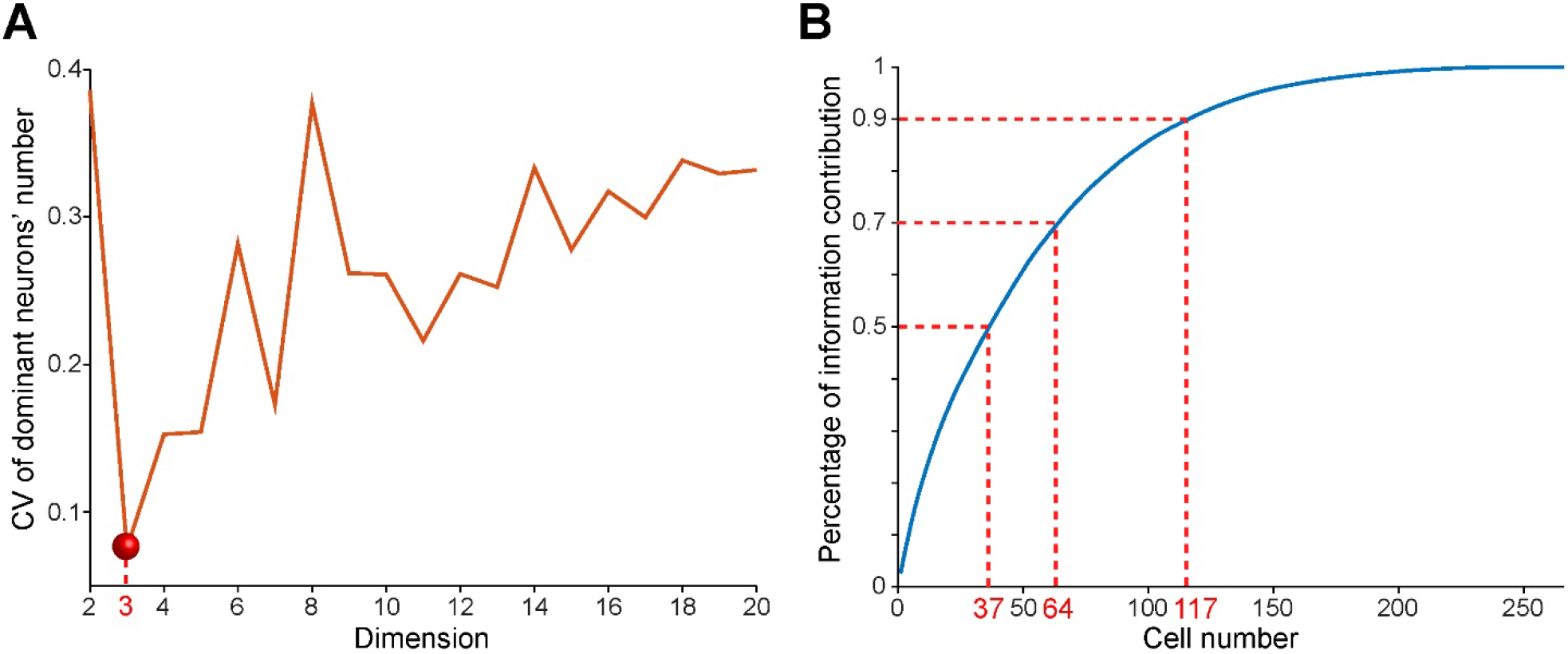
Selecting the best dimension and discovering cell assemblies. (A) Shown is an example of selecting the optimum dimension of ensemble patterns (sleep dataset). {*CV*_*i*_} of the number of large-contributed neurons are calculated for dimensions from 2 to 20, dimension 3 (highlighted by the red circle) with a minimum *CV* is selected as the best relevant dimension. (B) The blue curve shows the relationship between the number of neurons (x-axis) and their percentage of summing weights in the whole neural circuit (y-axis). 37 neurons (13.9% of total 266 neurons) contribute 50% information of corresponding ensemble pattern, 64 neurons (24.1%) contain 70% information contribution, while 90% of the information is contributed by 117 neurons (44.0%).

**Figure S4.**
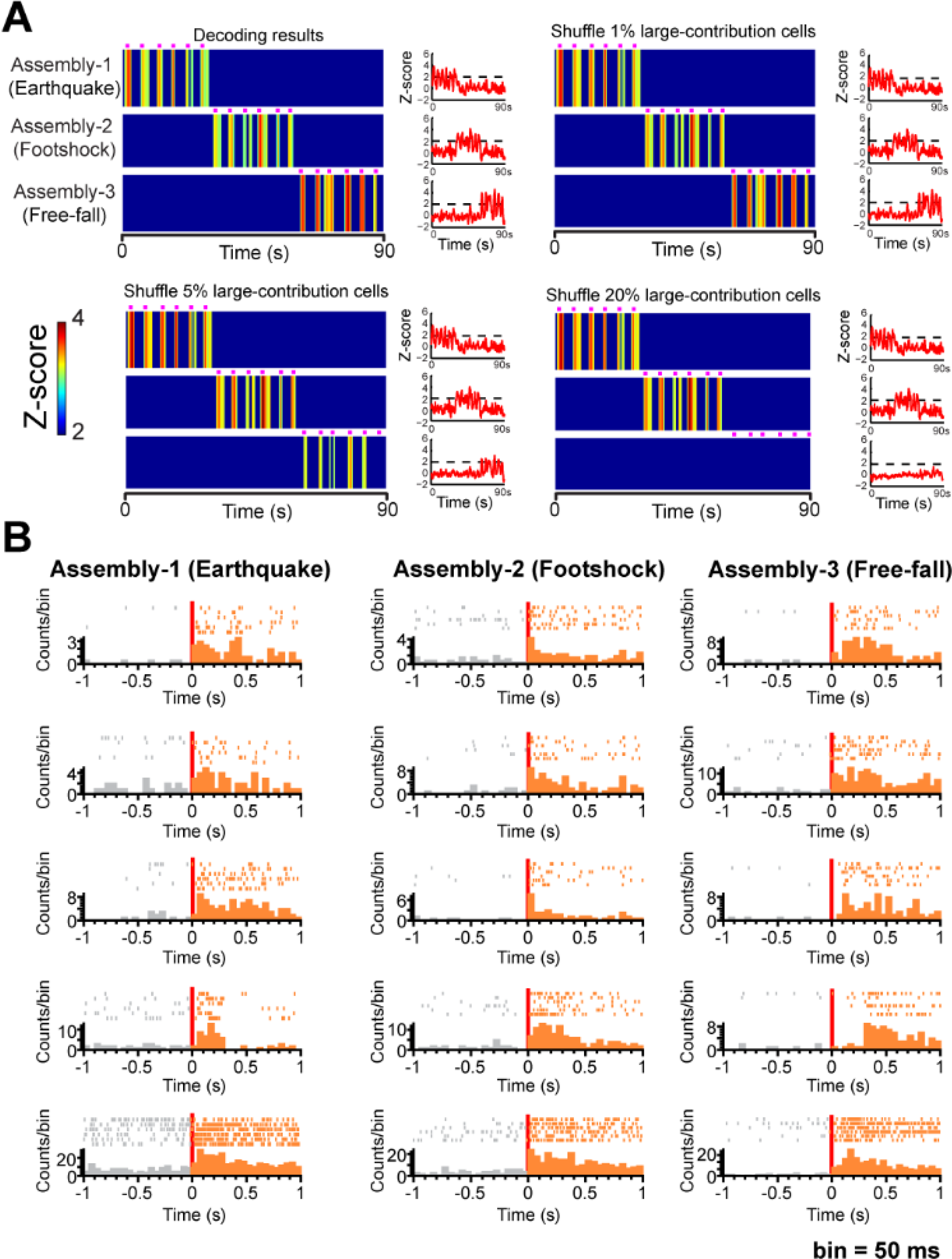
Verification of decoded cell assemblies by shuffling top-weight neurons and responsiveness of cell assemblies encoding three fearful experiences. (A) We shuffled top large-weight neurons by replacing their firing pattern with Gaussian noise with the same mean firing rate and standard deviation. We observed that the ensemble pattern of Assembly-2 (corresponding to Footshock) gradually becomes weaker as more top contributed neurons are shuffled, while leaving other two ensemble patterns representing the Earthquake or Free-fall unchanged (Six trials per event, as shown by six purple bars above each ensemble pattern). Ensemble pattern of Assembly-2 finally disappears when top 20% contributed cells are shuffled. (B) From left to right, each column shows peri-event rasters/histograms of five cell members of each cell assembly in response to the corresponding fearful episodic stimulus.

**Figure S5.**
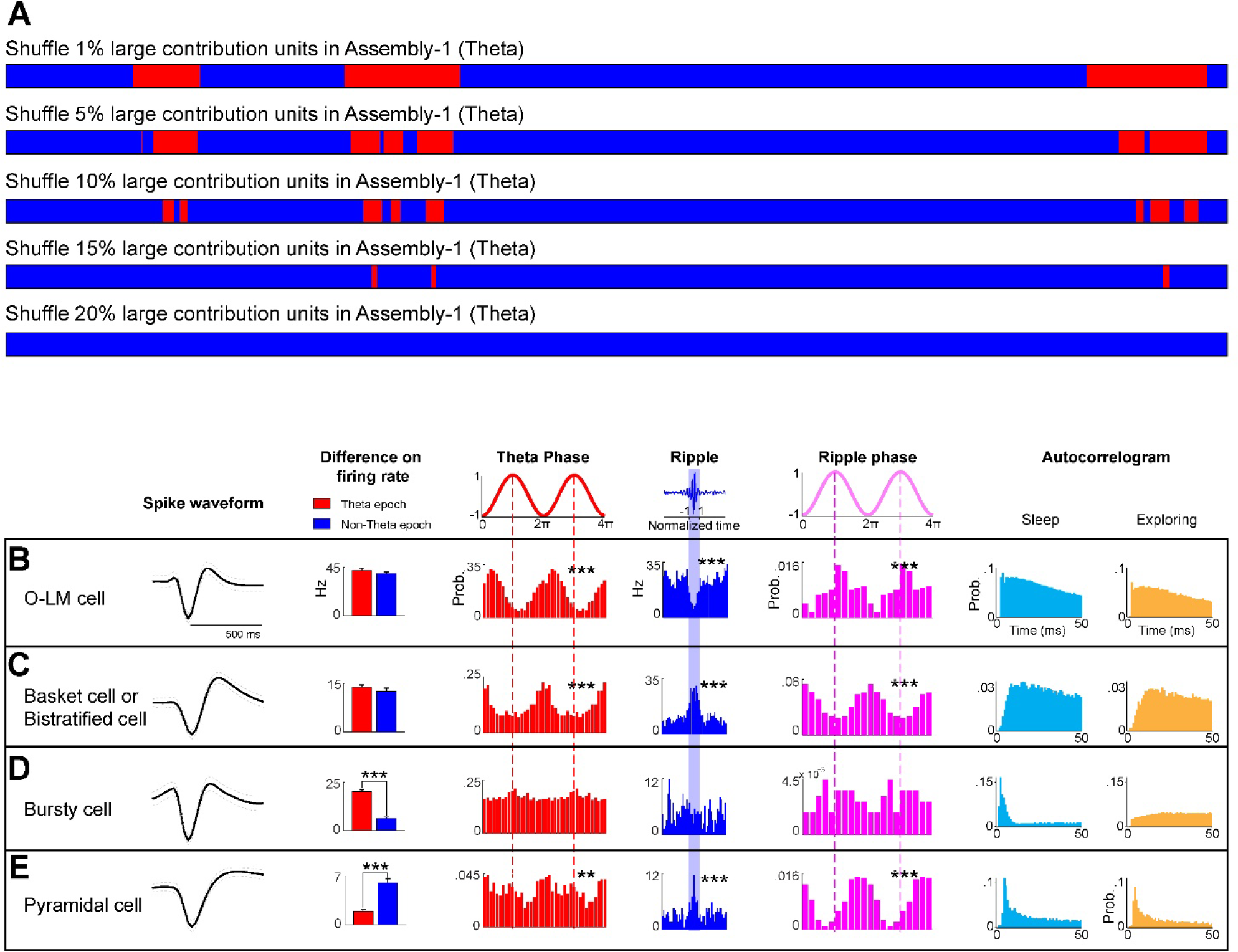
Verification of decoded cell assemblies by shuffling top-weight neurons and cell-type analysis. (A) Top-weight neurons were shuffled by replacing their firing pattern with Gaussian noise with the same mean firing rate and standard deviation. The ensemble pattern of Assembly-1 (corresponding to Theta oscillation) abate gradually when top contributed cells are shuffled. Furthermore, the ensemble pattern of Assembly-1 disappears when top 20% contributed cells are shuffled. (B-E) Firing properties of different cell types in cell assemblies related to sleep oscillations. Shown are examples of four putative cell types [O-LM cell (B), Basket/Bistratified cell (C), Bursty cell (D) and pyramidal cell (E)] identified based on their distinct firing properties. The 1st left subpanels of B-E show the average waveforms of representative units of four cell types. The 2nd left subpanels of B-E are the firing rate differences between Theta and non-Theta epoch. The 3rd left subpanels of B-E show their theta phase-coupling properties, and 4th and 5th subpanels of B-E are the firing properties and phase-coupling properties at ripple occurrences. The right two subpanels of B-E are auto-correlograms of spike activities during sleep and exploring states. Note the peaks within 10 ms in the auto-correlogram plots indicate burst firing. These four cell types show distinct firing properties. Specifically, O-LM cells (B) exhibit inhibited firing upon ripple occurrences, which is a peculiar feature among all these cell types. Basket/Bistratified cells (C) exhibit high firing rates upon ripples, these cells show strongly coupled spike timing of with Theta oscillations and ripples (P<0.001, Rayleigh’s test), and these cells fire at the troughs of Theta oscillations and ripples. The distinguishing features of Bursty cells (D) are twofold: significant higher firing rates within Theta epochs and bursty firing during sleep. Pyramidal cells (E) have wider spike waveforms and high probabilities of burst firing. Here, the firing rate differences are examined by *t*-test, and Rayleigh’s test is applied to phase-coupling properties of Theta oscillation and ripples. Two asterisks denote that P-value is in the range of 0.01-0.001, three asterisks denote P<0.001.

**Figure S6.**
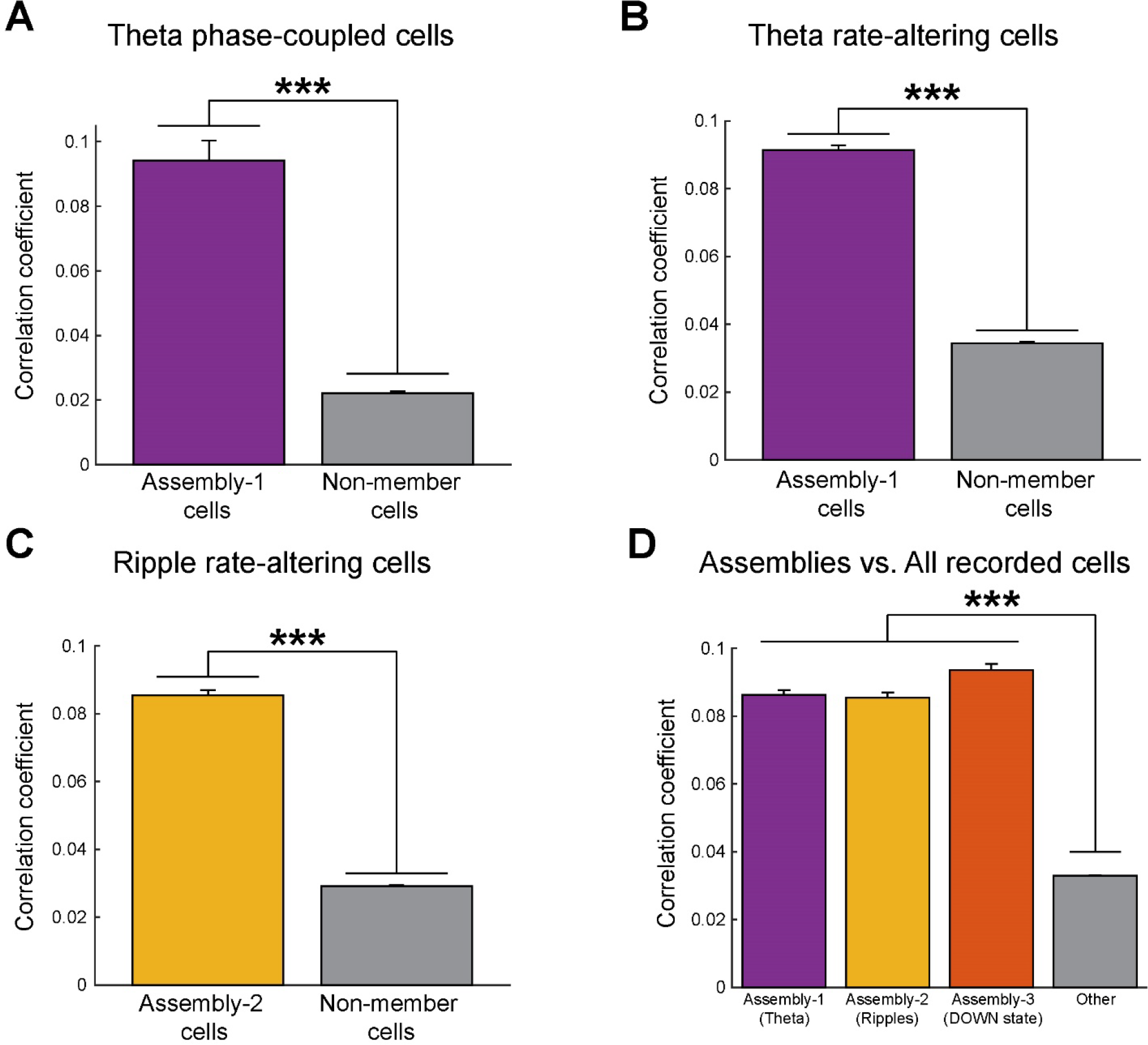
Correlation analysis further verified the cell assemblies related to sleep oscillations. We carry out correlation analysis on three cell assemblies corresponding to different sleep oscillations as well as other units. The neural data from the individual unit is first converted into a spike occurrence vector with a 200-ms bin size. We calculate Pearson’s correlation coefficients between the occurrence vectors of all simultaneously recorded neurons during animal sleeping. Correlation analysis is performed using the following formula:

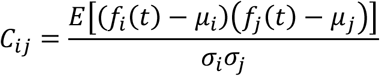 Where *f*_*i*_ (*t*) and *f*_*j*_ (*t*) are the two spike occurrence vectors converted from neural data of unit *i* and unit *j* simultaneously recorded from the same animal; and *μ*_*i*_ and *μ*_*j*_ are the mean values of these two vectors respectively, *σ*_*i*_ and *σ*_*j*_ are standard deviations. (A) There are 52 neurons show significant phase-coupling to Theta oscillation (Rayleigh’s circular statistics, p<0.001). Among these 52 neurons, 9 neurons are cell members of Assembly-1 (Theta), other 43 neurons aren’t. We calculate the correlations within these member and non-member theta phase-coupling cells, and observe that the correlations within Assembly-1 theta phase-coupling cells are significantly higher than the correlations within non-member theta phase-coupling cells (t-test: p<0.001. 0.0942± 0.0062 vs. 0.0221± 0.0006, data shown as mean±SE). (B) There are 150 cells exhibit a significant difference in averaged firing rates between theta epochs vs. non-theta epochs (t-test: p<0.001). 48 of these Theta rate-altering cells are cell members of Assembly-1 (Theta), and 102 cells aren’t. We observe that the correlations within Assembly-1 theta rate-altering cells are significantly higher than the correlations within non-member theta rate-altering cells (t-test: p<0.001. 0.0913± 0.0014 vs. 0.0344± 0.0004). (C) There are 239 cells exhibit significant firing rate changes during ripples (t-test: p<0.001). 53 of these ripple cells are the members of Assembly-2 (Ripples). The correlations within Assembly-2 cells are significantly higher than the correlations within non-member ripple cells (t-test: p<0.001.0 0854± 0.0015 vs. 0.0292± 0.0002). (D) Correlations within three cell assemblies (53 neurons in each cell assembly (top 20% of 266 neurons)) are significantly higher than correlations within the neural population (all 266 neurons) (one-way ANOVA with post-hoc test. Assembly-1: 0.0863± 0.0014, Assembly-2: 0.0854± 0.0015, Assembly-3: 0.0936± 0.0018, All: 0.0329± 0.0002).

**Figure S7.**
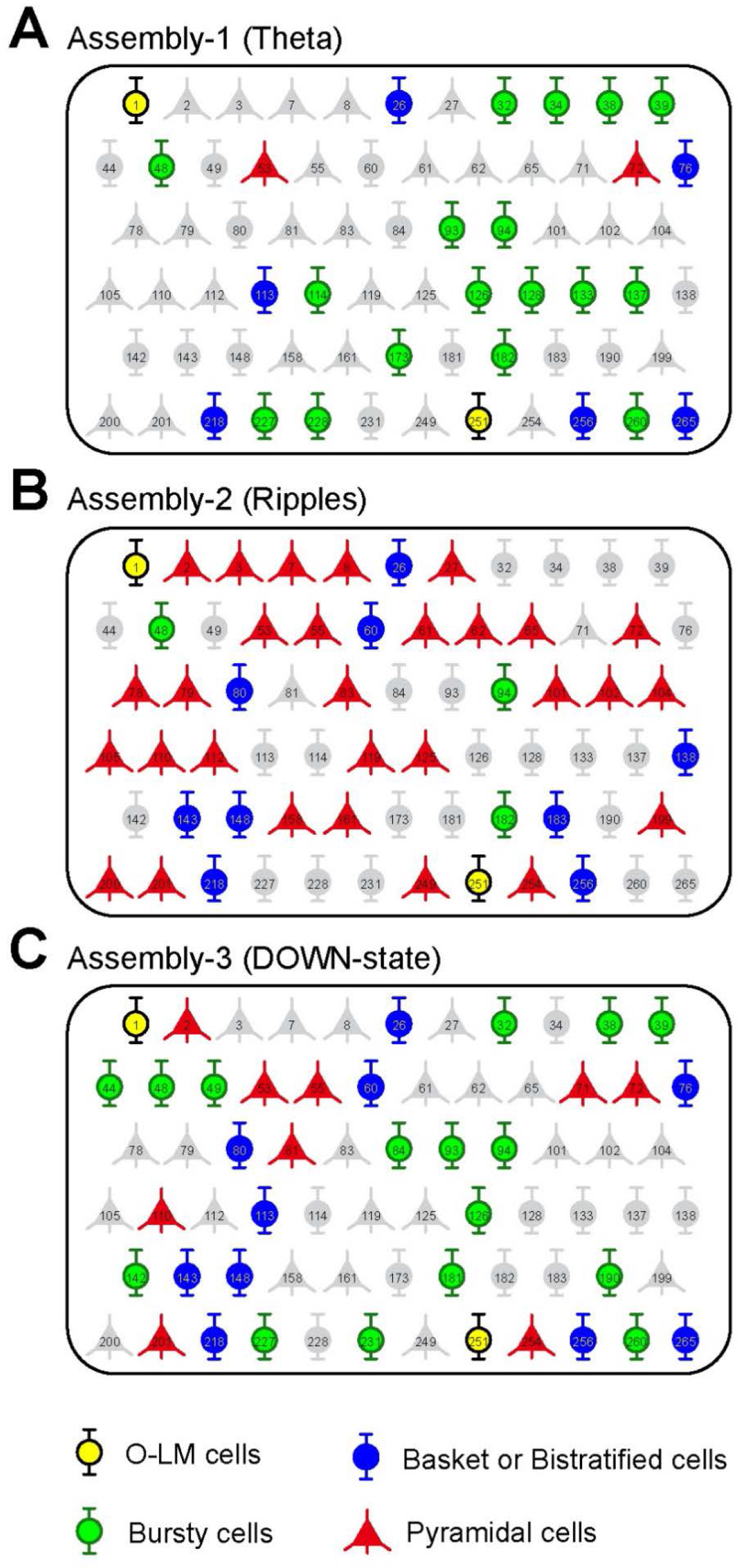
Combinatorial activation patterns among three distinct cell assemblies during animal’s sleep. (A) Illustration of identified Assembly-1 member cells engaged in sleep theta (in color: yellow for putative O-LM cells; blue for putative basket/bistratified interneurons; green for putative bursty interneurons; and red for putative pyramidal cells). Non-member cells are in gray. Numerical numbers inside each cell represent the unit simultaneously recorded. (B) Assembly-2 corresponding to ripples. (C) Assembly-3 corresponding to DOWN-state. Overall, these CA1 cell assemblies employed specific-to-general combinatorial coding strategy to represent each sleep oscillation. For example, two O-LM cells (unit-1 and unit-251) were identified in all three assemblies, so were some of the pyramidal cells (unit-53 and unit-72). Some of the pyramidal cells (unit-110 and unit-201) and basket/bistratified cells (i.e. unit-60, unit-80, etc.) participated in both Assembly-2 (ripples) and Assembly-3 (DOWN-state), whereas some of the bursty cells (i.e. unit-32, unit-38, unit-39, etc.) participated in Assembly-1 and Assembly-3. Many cells exhibited assembly-specific memberships. For instance, pyramidal cells unit-101, unit-102, and unit-200 were Assembly-2 specific; bursty cells unit-173 and unit-228 were specific members of Assembly-1.

**Figure S8.**
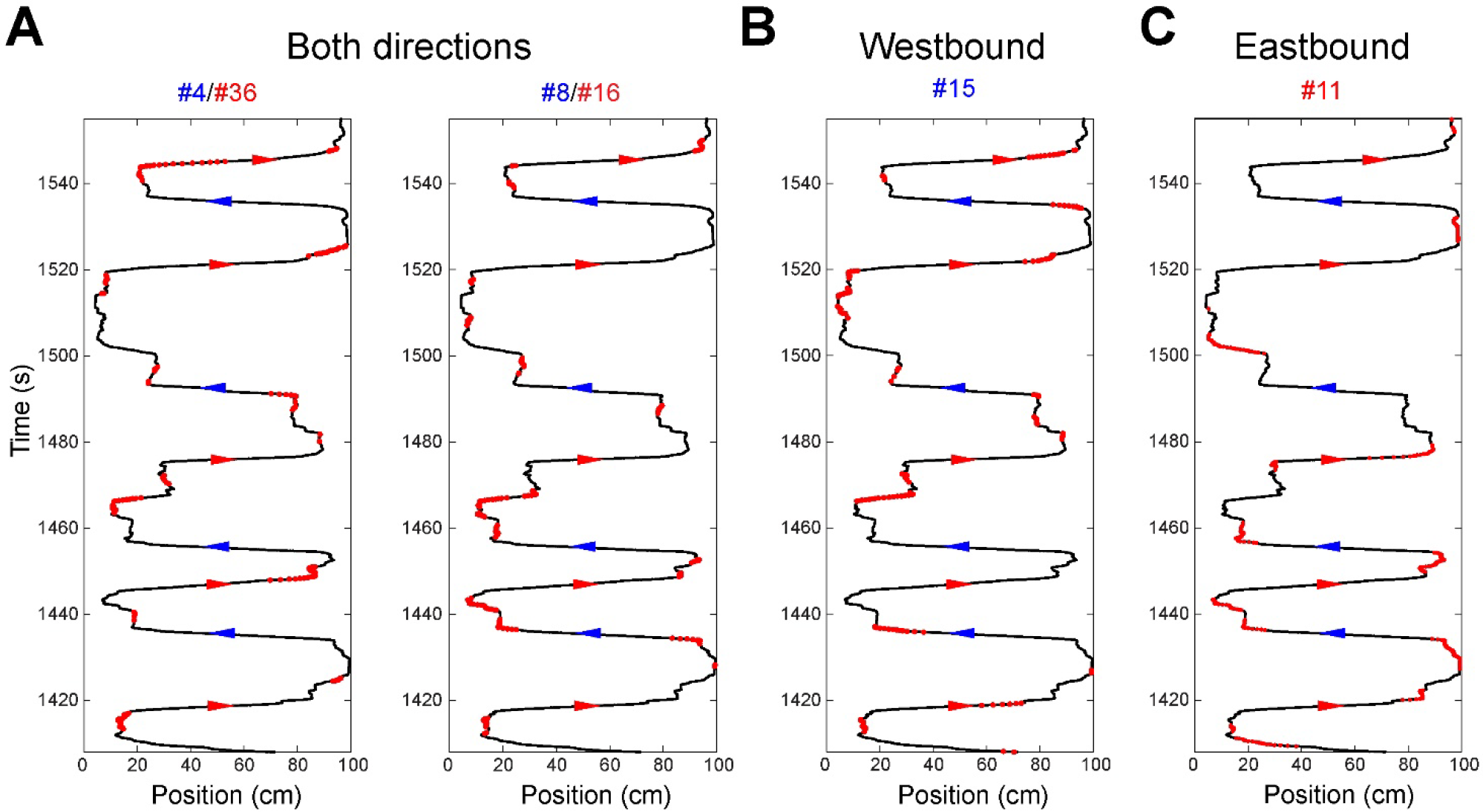
Start & finish cells in navigational experiences. Shown are the firing-location plots of four start & finish cells decoded from linear track experiment. All these cells exhibit higher firing rate at two ends of the linear track and decreased firing when animal running in the linear track. (A) shown are two start & finish cells participated in both cell assemblies. (B) Firing-location plot of a start & finish cell in Assembly-1 (Westbound), this cell is not an Assembly-2 (Eastbound) member. (C) An Assembly-2 (Eastbound) start & finish cell. Numbers above plots are their rankings among corresponding cell assemblies.

**Figure S9.**
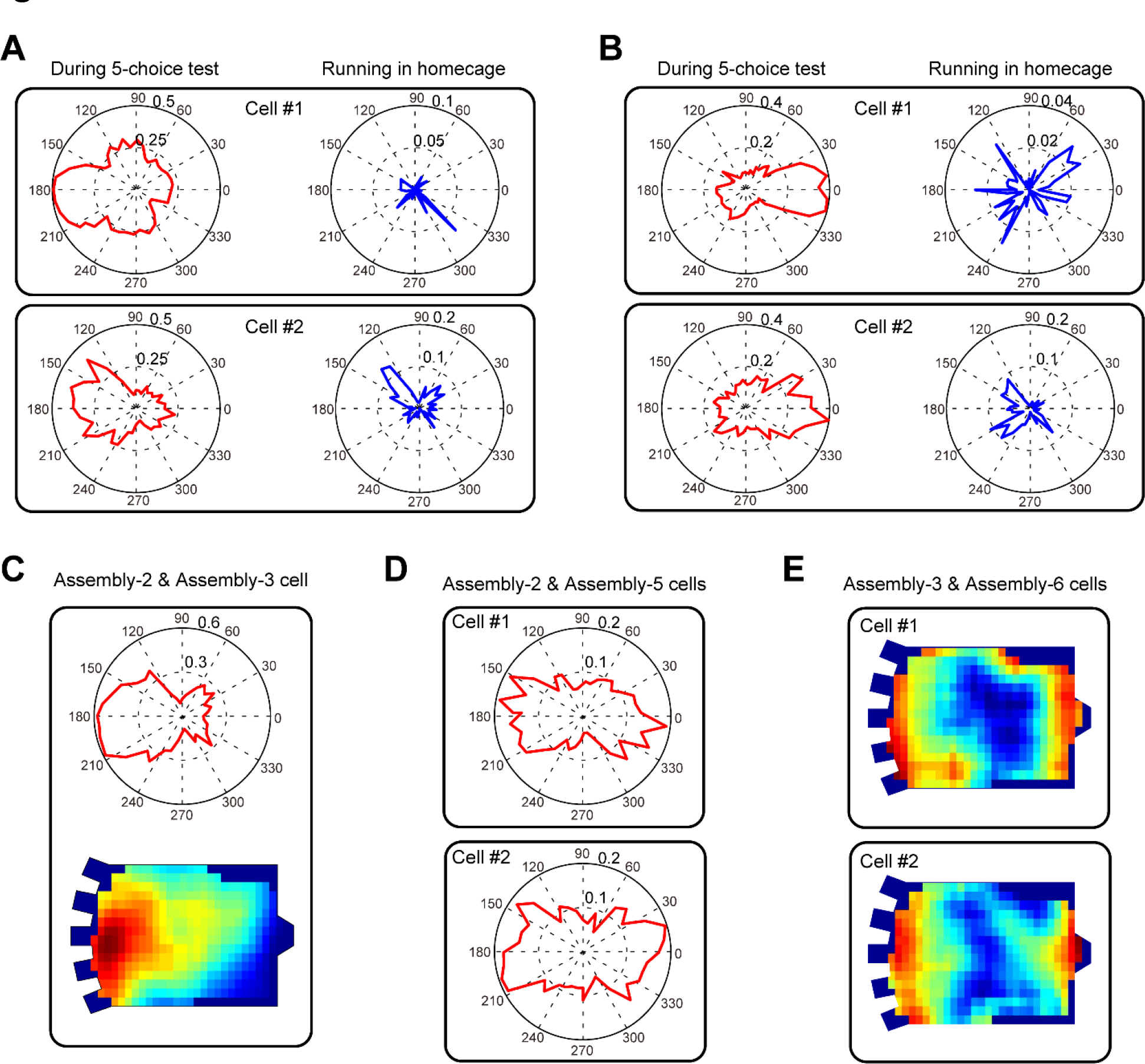
“Running” and “Returning” cells were task-specific as they showed no directional preferences in the home-cage. (A) Two representative units of “Running” cell assembly. Red curves are orientation-firing polar histograms during five-choice discrimination operant conditioning task, blue curves are orientation-firing polar histograms when animal running (speed ≥2.5cm/s) in its home-cage. (B) Two representative units of “Returning” cell assembly. It can be observed that units in a and b have no direction-specific firing when the animal in its home-cage environment. **Cells participated in multiple cell assemblies.** (C) Orientation-firing polar histogram and location-firing map of a neuron participated in Assembly-2 (Running) and Assembly-3 (Poke zone). This neuron shows both orientation-and location-firing properties which is consistent with the definitions of ensemble patterns of “Running” and “Poke zone”. (D) Orientation-firing polar histograms of two cells participated in Assembly-2 (Running) and Assembly-5 (Returning). These two cells show orientation-firing on both directions. (E) Location-firing maps of two cells participated in Assembly-3 (Poke zone) and Assembly-6 (Reward zone). Location-firings of these two cells are significantly higher on both sides. The dotted cycles and the numbers in the orientation-firing polar histograms of the cells in (A-D) denote the probabilities of neuronal spike firing in the corresponding orientation.

**Figure S10.**
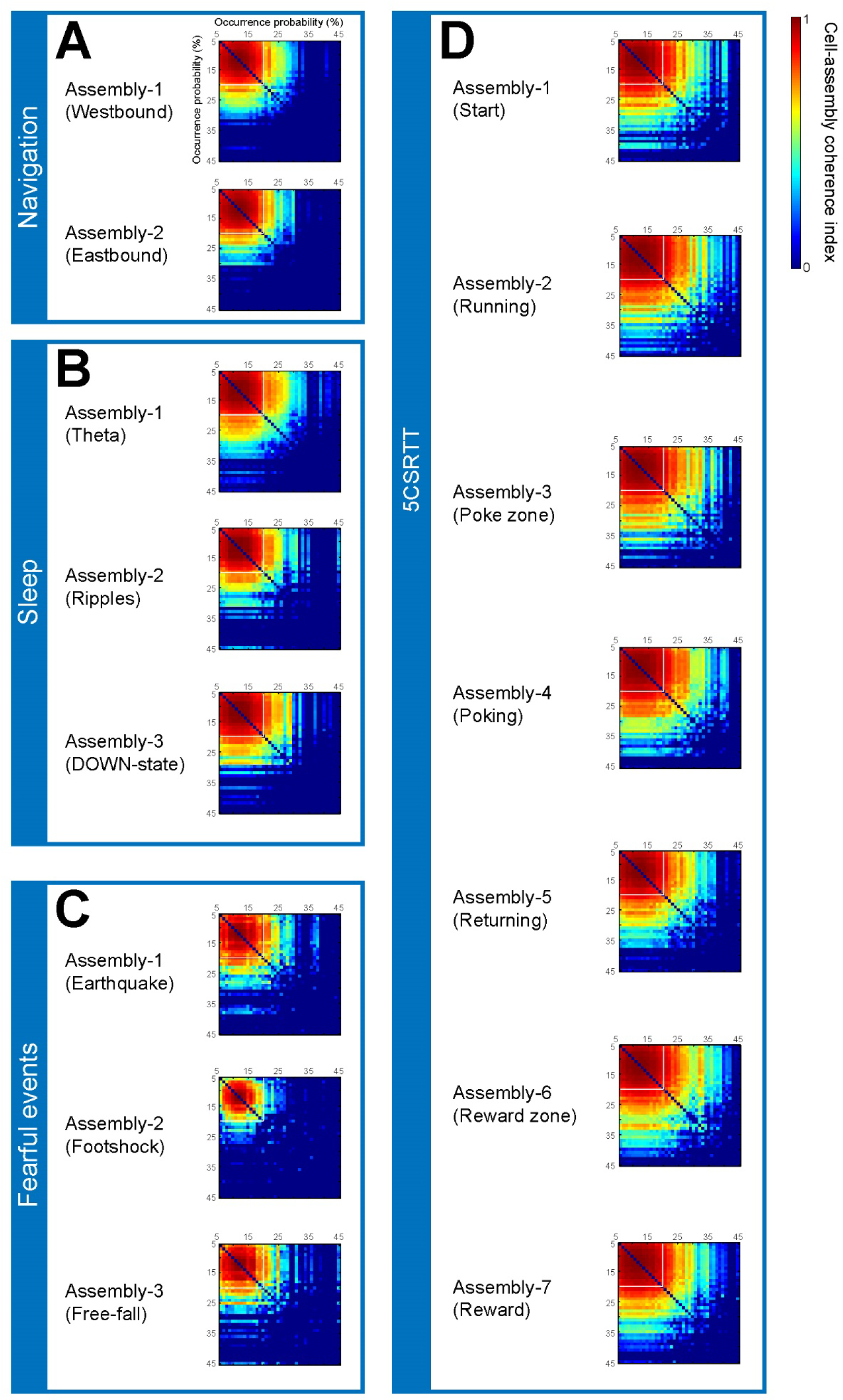
We systemically vary lower thresholds from 5% to 45% with step by 1% (the corresponding upper thresholds are set as one minus the value of lower point accordingly) and calculate the cell-assembly coherence indexes between the ensemble patterns corresponding to same state/stimuli. Shown are cell-assembly coherence matrixes of all 15 cell assemblies in four different experiments (A-D), it is thus evident that robust ensemble patterns can be obtained using 5%-20% ranges.

## References

1. Adrian ED. The basis of sensation. 1928.

2. Faisal AA, Selen LP, Wolpert DM. Noise in the nervous system. Nat Rev Neurosci. 2008;9(4):292–303.

3. Rolls E, Deco G. The noisy brain. Stochastic dynamics as a principle of brain function(Oxford Univ Press, UK, 2010). 2010.

4. Shadlen MN, Newsome WT. Noise, neural codes and cortical organization. Curr Opin Neurobiol. 1994;4(4):569–79.

5. Stein RB, Gossen ER, Jones KE. Neuronal variability: noise or part of the signal? Nat Rev Neurosci. 2005;6(5):389–97.

6. Averbeck BB, Latham PE, Pouget A. Neural correlations, population coding and computation. Nat Rev Neurosci. 2006;7(5):358–66.

7. Kanitscheider I, Coen-Cagli R, Pouget A. Origin of information-limiting noise correlations. Proc Natl Acad Sci U S A. 2015;112(50):E6973–82.

8. Lee D, Port NL, Kruse W, Georgopoulos AP. Variability and correlated noise in the discharge of neurons in motor and parietal areas of the primate cortex. J Neurosci. 1998;18(3):1161–70.

9. Hartmann C, Lazar A, Nessler B, Triesch J. Where’s the Noise? Key Features of Spontaneous Activity and Neural Variability Arise through Learning in a Deterministic Network. PLoS Comput Biol. 2015;11(12):e1004640.

10. Kohn A, Coen-Cagli R, Kanitscheider I, Pouget A. Correlations and Neuronal Population Information. Annu Rev Neurosci. 2016;39:237–56.

11. Abbott L, Sejnowski TJ. Neural codes and distributed representations: foundations of neural computation: Mit Press; 1999.

12. Mainen ZF, Sejnowski TJ. Reliability of spike timing in neocortical neurons. Science. 1995;268(5216):1503–6.

13. Abbott LF, Dayan P. The effect of correlated variability on the accuracy of a population code. Neural Comput. 1999; 11(1):91–101.

14. Arieli A, Sterkin A, Grinvald A, Aertsen A. Dynamics of ongoing activity: explanation of the large variability in evoked cortical responses. Science. 1996;273(5283): 1868–71.

15. Churchland MM, Yu BM, Cunningham JP, Sugrue LP, Cohen MR, Corrado GS, et al. Stimulus onset quenches neural variability: a widespread cortical phenomenon. Nat Neurosci. 2010;13(3):369–78.

16. Lin IC, Okun M, Carandini M, Harris KD. The Nature of Shared Cortical Variability. Neuron. 2015;87(3):644–56.

17. Luczak A, Bartho P, Harris KD. Gating of sensory input by spontaneous cortical activity. The Journal of neuroscience: the official journal of the Society for Neuroscience. 2013;33(4):1684–95.

18. Marcos E, Pani P, Brunamonti E, Deco G, Ferraina S, Verschure P. Neural variability in premotor cortex is modulated by trial history and predicts behavioral performance. Neuron. 2013;78(2):249–55.

19. Petersen CCH, Hahn TTG, Mehta M, Grinvald A, Sakmann B. Interaction of sensory responses with spontaneous depolarization in layer 2/3 barrel cortex. P Natl Acad Sci USA. 2003; 100(23):13638–43.

20. Saberi-Moghadam S, Ferrari-Toniolo S, Ferraina S, Caminiti R, Battaglia-Mayer A. Modulation of Neural Variability in Premotor, Motor, and Posterior Parietal Cortex during Change of Motor Intention. The Journal of neuroscience: the official journal of the Society for Neuroscience. 2016;36(16):4614–23.

21. Scholvinck ML, Saleem AB, Benucci A, Harris KD, Carandini M. Cortical state determines global variability and correlations in visual cortex. The Journal of neuroscience: the official journal of the Society for Neuroscience. 2015;35(1):170–8.

22. Wehr M, Zador AM. Balanced inhibition underlies tuning and sharpens spike timing in auditory cortex. Nature. 2003;426(6965):442–6.

23. Fenton AA, Muller RU. Place cell discharge is extremely variable during individual passes of the rat through the firing field. P Natl Acad Sci USA. 1998;95(6):3182–7.

24. Johnson A, Fenton AA, Kentros C, Redish AD. Looking for cognition in the structure within the noise. Trends Cogn Sci. 2009;13(2):55–64.

25. Lin L, Osan R, Tsien JZ. Organizing principles of real-time memory encoding: neural clique assemblies and universal neural codes. Trends Neurosci. 2006;29(1):48–57.

26. Li M, Tsien J. Neural Code-Neural Self-Information Theory on How Cell-Assembly Code Rises from Spike Time and Neuronal Variability. Frontiers in Cellular Neuroscience. 2017;11:236.

27. Hebb DO. The organization of behavior: A neuropsychological approach: John Wiley & Sons; 1949.

28. Buzsaki G. Neural syntax: cell assemblies, synapsembles, and readers. Neuron. 2010;68(3):362–85.

29. Eggermont JJ. Is there a neural code? Neurosci Biobehav Rev. 1998;22(2):355–70.

30. Harris KD, Csicsvari J, Hirase H, Dragoi G, Buzsaki G. Organization of cell assemblies in the hippocampus. Nature. 2003;424(6948):552–6.

31. Maurer AP, Cowen SL, Burke SN, Barnes CA, McNaughton BL. Organization of hippocampal cell assemblies based on theta phase precession. Hippocampus. 2006; 16(9):785–94.

32. Palm G. Towards a theory of cell assemblies. Biol Cybern. 1981;39(3):181–94.

33. Maimon G, Assad JA. Beyond Poisson: increased spike-time regularity across primate parietal cortex. Neuron. 2009;62(3):426–40.

34. Xie K, Kuang H, Tsien JZ. Mild blast events alter anxiety, memory, and neural activity patterns in the anterior cingulate cortex. PLoS One. 2013;8(5):e64907.

35. Bliss TVP, Collingridge GL, Kaang B-K, Zhuo M. Synaptic plasticity in the anterior cingulate cortex in acute and chronic pain. Nat Rev Neurosci. 2016;17(8):485–96.

36. Steenland HW, Li X-Y, Zhuo M. Predicting aversive events and terminating fear in the mouse anterior cingulate cortex during trace fear conditioning. The Journal of neuroscience: the official journal of the Society for Neuroscience. 2012;32(3):1082–95.

37. Liu J, Wei W, Kuang H, Tsien JZ, Zhao F. Heart rate and heart rate variability assessment identifies individual differences in fear response magnitudes to earthquake, free fall, and air puff in mice. PLoS One. 2014;9(3):e93270.

38. Cantero JL, Atienza M, Stickgold R, Kahana MJ, Madsen JR, Kocsis B. Sleep-dependent theta oscillations in the human hippocampus and neocortex. The Journal of neuroscience: the official journal of the Society for Neuroscience. 2003;23(34):10897–903.

39. Colgin LL. Rhythms of the hippocampal network. Nat Rev Neurosci. 2016;17(4):239–49.

40. Csicsvari J, Hirase H, Czurko A, Mamiya A, Buzsaki G. Oscillatory coupling of hippocampal pyramidal cells and interneurons in the behaving Rat. The Journal of neuroscience: the official journal of the Society for Neuroscience. 1999;19(1):274–87.

41. Fox SE, Wolfson S, Ranck JBJr., Hippocampal theta rhythm and the firing of neurons in walking and urethane anesthetized rats. Exp Brain Res. 1986;62(3):495–508.

42. Chen G, Wang LP, Tsien JZ. Neural population-level memory traces in the mouse hippocampus. PLoS One. 2009;4(12):e8256.

43. Sara SJ. The locus coeruleus and noradrenergic modulation of cognition. Nat Rev Neurosci. 2009;10(3):211–23.

44. Zhang H, Chen G, Kuang H, Tsien JZ. Mapping and deciphering neural codes of NMDA receptor-dependent fear memory engrams in the hippocampus. PLoS One. 2013;8(11):e79454.

45. Klausberger T, Somogyi P. Neuronal diversity and temporal dynamics: the unity of hippocampal circuit operations. Science. 2008;321(5885):53–7.

46. Kuang H, Lin L, Tsien JZ. Temporal dynamics of distinct CA1 cell populations during unconscious state induced by ketamine. PLoS One. 2010;5(12):e15209.

47. Somogyi P, Klausberger T. Defined types of cortical interneurone structure space and spike timing in the hippocampus. J Physiol. 2005;562(Pt 1):9–26.

48. Diba K, Buzsaki G. Forward and reverse hippocampal place-cell sequences during ripples. Nat Neurosci. 2007;10(10):1241–2.

49. Dupret D, O’Neill J, Pleydell-Bouverie B, Csicsvari J. The reorganization and reactivation of hippocampal maps predict spatial memory performance. Nat Neurosci. 2010;13(8):995–1002.

50. Pastalkova E, Itskov V, Amarasingham A, Buzsaki G. Internally generated cell assembly sequences in the rat hippocampus. Science. 2008;321(5894):1322–7.

51. Trimper JB, Trettel SG, Hwaun E, Colgin LL. Methodological Caveats in the Detection of Coordinated Replay between Place Cells and Grid Cells. Front Syst Neurosci. 2017;11.

52. Wilson MA, McNaughton BL. Reactivation of hippocampal ensemble memories during sleep. Science. 1994;265(5172):676–9.

53. Gupta AS, van der Meer MA, Touretzky DS, Redish AD. Hippocampal replay is not a simple function of experience.Neuron. 2010;65(5):695–705.

54. Kelemen E, Fenton AA. Coordinating different representations in the hippocampus. Neurobiol Learn Mem. 2016;129:50–9.

55. Brown EN, Kass RE, Mitra PP. Multiple neural spike train data analysis: state-of-the-art and future challenges. Nat Neurosci. 2004;7(5):456–61.

56. Ermentrout GB, Galan RF, Urban NN. Reliability, synchrony and noise. Trends Neurosci. 2008;31(8):428–34.

57. Masquelier T. Neural variability, or lack thereof. Front Comput Neurosci. 2013;7:7.

58. Giustino TF, Fitzgerald PJ, Maren S. Fear Expression Suppresses Medial Prefrontal Cortical Firing in Rats. PloS one. 2016;11(10):e0165256.

59. Kim JJ, Jung MW. Neural circuits and mechanisms involved in Pavlovian fear conditioning: a critical review. Neurosci Biobehav Rev. 2006;30(2):188–202.

60. Zhuo M. Neural Mechanisms Underlying Anxiety-Chronic Pain Interactions. Trends Neurosci. 2016;39(3):136–45.

61. Tsien JZ. A Postulate on the Brain’s Basic Wiring Logic. Trends Neurosci. 2015;38(11):669–71.

62. Xie K, Fox GE, Liu J, Lyu C, Lee JC, Kuang H, et al. Brain Computation Is Organized via Power-of-Two-Based Permutation Logic. Front Syst Neurosci. 2016;10:95.

63. Fenton AA, Kao H-Y, Neymotin SA, Olypher A, Vayntrub Y, Lytton WW, et al. Unmasking the CA1 ensemble place code by exposures to small and large environments: more place cells and multiple, irregularly arranged, and expanded place fields in the larger space. The Journal of neuroscience: the official journal of the Society for Neuroscience. 2008;28(44):11250–62.

64. Park E, Dvorak D, Fenton AA. Ensemble place codes in hippocampus: CA1, CA3, and dentate gyrus place cells have multiple place fields in large environments. PloS one. 2011;6(7):e22349.

65. Hok V, Lenck-Santini P-P, Roux S, Save E, Muller RU, Poucet B. Goal-related activity in hippocampal place cells. The Journal of neuroscience: the official journal of the Society for Neuroscience. 2007;27(3):472–82.

66. Ito HT, Zhang S-J, Witter MP, Moser EI, Moser M-B. A prefrontal-thalamo-hippocampal circuit for goal-directed spatial navigation. Nature. 2015;522(7554):50–5.

67. Wikenheiser AM, Redish AD. Changes in reward contingency modulate the trial-to-trial variability of hippocampal place cells. J Neurophysiol. 2011;106(2):589–98.

68. Eschenko O, Ramadan W, Molle M, Born J, Sara SJ. Sustained increase in hippocampal sharp-wave ripple activity during slow-wave sleep after learning. Learn Mem. 2008;15(4):222–8.

69. Sara SJ. Sleep to Remember. The Journal of neuroscience: the official journal of the Society for Neuroscience. 2017;37(3):457–63.

70. Graves AR, Moore SJ, Bloss EB, Mensh BD, Kath WL, Spruston N. Hippocampal pyramidal neurons comprise two distinct cell types that are countermodulated by metabotropic receptors. Neuron. 2012;76(4):776–89.

71. Mizuseki K, Diba K, Pastalkova E, Buzsaki G. Hippocampal CA1 pyramidal cells form functionally distinct sublayers. Nat Neurosci. 2011;14(9): 1174–81.

72. Tsien JZ. Cre-Lox Neurogenetics: 20 Years of Versatile Applications in Brain Research and Counting. Front Genet. 2016;7:19.

73. Gan J, Weng S-M, Pernia-Andrade AJ, Csicsvari J, Jonas P. Phase-Locked Inhibition, but Not Excitation, Underlies Hippocampal Ripple Oscillations in Awake Mice InVivo.Neuron. 2017;93(2):308–14.

74. Li Y, Xu J, Liu Y, Zhu J, Liu N, Zeng W, et al. A distinct entorhinal cortex to hippocampal CA1 direct circuit for olfactory associative learning. Nat Neurosci. 2017;20(4):559–70.

75. Nagode DA, Tang A-H, Yang K, Alger BE. Optogenetic identification of an intrinsic cholinergically driven inhibitory oscillator sensitive to cannabinoids and opioids in hippocampal CA1. J Physiol. 2014;592(1):103–23.

76. Bari A, Dalley JW, Robbins TW. The application of the 5-choice serial reaction time task for the assessment of visual attentional processes and impulse control in rats. Nat Protoc. 2008;3(5):759–67.

77. Kim H, Ahrlund-Richter S, Wang X, Deisseroth K, Carlen M. Prefrontal Parvalbumin Neurons in Control of Attention. Cell. 2016;164(1-2):208–18.

78. Totah NKB, Kim YB, Homayoun H, Moghaddam B. Anterior cingulate neurons represent errors and preparatory attention within the same behavioral sequence. The Journal of neuroscience: the official journal of the Society for Neuroscience. 2009;29(20):6418–26.

79. Yin HH. Action, time and the basal ganglia. Philos Trans R Soc Lond B Biol Sci. 2014;369(1637):20120473.

80. Laubach M. A comparative perspective on executive and motivational control by the medial prefrontal cortex. Neural Basis of Motivational and Cognitive Control. 2011:95.

81. Miller EK. The prefrontal cortex and cognitive control. Nat Rev Neurosci. 2000;1(1):59–65.

82. Giustino TF, Maren S. The Role of the Medial Prefrontal Cortex in the Conditioning and Extinction of Fear. Front Behav Neurosci. 2015;9:298.

83. Hardy M. Pareto’s law. The Mathematical Intelligencer. 2010;32(3):38–43.

84. Iqbal M, Rizwan M, editors. Application of 80/20 rule in software engineering Waterfall Model. Information and Communication Technologies, 2009 ICICT’09 International Conference on; 2009: IEEE.

85. Juran JM. Pareto, lorenz, cournot, bernoulli, juran and others. 1950.

86. Ultsch A. Proof of Pareto’s 80/20 law and Precise Limits for ABC-Analysis. Data Bionics Research Group University of Marburg/Lahn, Germany. 2002:1–11.

87. Lin L, Chen G, Xie K, Zaia KA, Zhang S, Tsien JZ. Large-scale neural ensemble recording in the brains of freely behaving mice. J Neurosci Methods. 2006;155(1):28–38.

88. Schmitzer-Torbert N, Jackson J, Henze D, Harris K, Redish AD. Quantitative measures of cluster quality for use in extracellular recordings. Neuroscience. 2005; 131(1):1–11.

89. Liu J, Wei W, Kuang H, Zhao F, Tsien JZ. Changes in heart rate variability are associated with expression of short-term and long-term contextual and cued fear memories. PLoS One. 2013;8(5):e63590.

90. Buzsaki G, Buhl DL, Harris KD, Csicsvari J, Czeh B, Morozov A. Hippocampal network patterns of activity in the mouse. Neuroscience. 2003; 116(1):201–11.

## References

1. Lin L, etal. (2006) Large-scale neural ensemble recording in the brains of freely behaving mice. J Neurosci Methods 155(1):28–38.

2. Xie K, et al. (2016) Brain Computation Is Organized via Power-of-Two-Based Permutation Logic. Front Syst Neurosci 10:95.

3. Paxinos G & Franklin KB (2004) The mouse brain in stereotaxic coordinates (Gulf Professional Publishing).

4. Kuang H, Lin L, & Tsien JZ (2010) Temporal dynamics of distinct CA1 cell populations during unconscious state induced by ketamine. PLoS One 5(12):e15209.

5. Schmitzer-Torbert N, Jackson J, Henze D, Harris K, & Redish AD (2005) Quantitative measures of cluster quality for use in extracellular recordings. Neuroscience 131(1):1–11.

6. Maimon G & Assad JA (2009) Beyond Poisson: increased spike-time regularity across primate parietal cortex. Neuron 62(3):426–440.

7. Hyvarinen A (1999) Fast and robust fixed-point algorithms for independent component analysis. IEEE Trans Neural Netw 10(3):626–634.

8. Liu J, Wei W, Kuang H, Tsien JZ, & Zhao F (2014) Heart rate and heart rate variability assessment identifies individual differences in fear response magnitudes to earthquake, free fall, and air puff in mice. PLoS One 9(3):e93270.

9. Liu J, Wei W, Kuang H, Zhao F, & Tsien JZ (2013) Changes in heart rate variability are associated with expression of short-term and long-term contextual and cued fear memories. PLoS One 8(5):e63590.

10. Buzsaki G, et al. (2003) Hippocampal network patterns of activity in the mouse. Neuroscience 116(1):201–211.

